# STAT3 restricts prostate cancer metastasis and antiandrogen resistance by controlling LKB1/CREB signaling pathway

**DOI:** 10.1101/2022.08.25.504915

**Authors:** Jan Pencik, Cecile Philippe, Michaela Schlederer, Matteo Pecoraro, Sandra Grund-Gröschke, Wen Jess Li, Amanda Tracz, Isabel Heidegger, Sabine Lagger, Karolína Trachtová, Monika Oberhuber, Ellen Heitzer, Osman Aksoy, Heidi A. Neubauer, Bettina Wingelhofer, Anna Orlova, Nadine Witzeneder, Thomas Dillinger, Elisa Redl, Georg Greiner, David D’Andrea, Johnny R. Östman, Simone Tangermann, Ivana Hermanova, Georg Schäfer, Adam Varady, Jaqueline Horvath, Dagmar Stoiber, Timothy I. Malcolm, Suzanne D. Turner, Eileen Parkes, Brigitte Hantusch, Gerda Egger, Stefan Rose-John, Valeria Poli, Suneil Jain, Chris W.D. Armstrong, Gregor Hoermann, Vincent Goffin, Fritz Aberger, Richard Moriggl, Arkaitz Carracedo, Cathal McKinney, Richard D Kennedy, Helmut Klocker, Michael R. Speicher, Dean G. Tang, Matthias Mann, Ali A. Moazzami, David M. Heery, Marcus Hacker, Lukas Kenner

## Abstract

Prostate cancer (PCa) lethality is driven by its progression to a metastatic castration-resistant state, yet the signaling mechanisms underlying metastatic spread remain unknown. Here we show that STAT3 converges with the LKB1/mTORC1 and CREB to control metastatic disease in PCa mouse models. Unexpectedly, STAT3 was found to be upregulated in diabetic PCa patients undergoing metformin therapy with a concomitant reduction in mTORC1 expression. In preclinical mouse models of PCa, genetic ablation or activation of STAT3 had opposing effects on LKB1/AMPK/mTORC1- dependent tumorigenesis. Using genetic and pharmacological approaches, we identified LKB1 as a direct STAT3 target while repressing CREB. Furthermore, PCa patients with high CREB expression had inferior clinical outcome with significantly increased risk of disease and metastatic recurrence. We observe that castration state lowers STAT3 abundance and increases AR and CREB levels, leading to castration-resistant PCa (CRPC). Our findings revealed that STAT3 controls mTORC1 and CREB in metastatic disease, suggesting CREB as a promising target for lethal CRPC.

## Introduction

Metastatic prostate cancer (PCa) is one of the leading causes of cancer death in men worldwide^1^. Up to 20% of patients present with primary metastatic disease, for which androgen-deprivation therapy (ADT) in combination with either second-generation ADT (abiraterone, enzalutamide or apalutamide) or docetaxel chemotherapy are the first line therapies for metastatic disease^2^. Although PCa is androgen receptor (AR) and androgen dependent, other signaling pathways including JAK/STAT proteins converge to drive disease progression and therapy resistance. Signal transducer and activator of transcription 3 (STAT3) is a complex transcriptional regulator responsible for essential biological functions such as differentiation, cell proliferation, immune responses and metabolism^3^. STAT3 signaling has both tumor suppressive or oncogenic roles in specific tissue contexts and is implicated in the regulation of malignant transformation and metastatic dissemination. For example, loss of STAT3 expression was found to synergize with driver mutations to promote brain^4^ or melanoma metastasis^5^, whereas, in lung cancer and PCa mouse models disruption of STAT3 suppresses lung or prostate metastasis compared to the wild-type cohort^6–8^. STAT3 transcriptional activity can trigger a metabolic switch toward aerobic glycolysis and regulate mitochondrial activity, a prominent metabolic feature of tumor cells^9–11^. The serine/threonine kinase mTOR is a master regulator of cell growth and metabolism and it can be negatively regulated by LKB1 (also known as serine/threonine kinase 11 or STK11)^12^. Activation of mTOR leads to the phosphorylation of eukaryotic initiation factor 4E-binding protein-1 (4E-BP1) and promotes protein synthesis^13^. mTOR promotes the phosphorylation of hundreds of substrates directly or indirectly via activating downstream kinases, including ribosomal

S6 kinase (S6K), and subsequent phosphorylation of S6 ribosomal protein to stimulate protein translation and cell proliferation. mTOR signaling is also frequently activated in cancer and is associated with various diseases such as obesity or type 2 diabetes mellitus (t2DM)^14^. A wide range of mTORC1 inhibitors have shown antitumor activity in glioblastoma^15^ and PCa patients^16^. PI3K/mTOR inhibition activates AR signaling in human xenograft and transgenic mouse models of PCa^17^. However, the emergence of resistance to mTORC1 inhibitors efficacy resulted in failure to improve patient outcomes in clinical trials to date^18^. Several factors might explain this limited impact in clinical applications, including heterogeneous intratumoral mTORC1 activity, resistant mutations of mTOR and activation of alternative proliferative signaling pathways^19^. Metformin, a first-line treatment for patients with t2DM, activates AMP-activated protein kinase (AMPK), which results in inhibition of mTORC1 in hepatocytes^20^. Metformin inhibits glucose and glutamine utilization in mitochondria and has been associated with decreased cancer risk in epidemiological studies in t2DM patients^21^ and in a variety of diabetic animal models^22^. Large-scale observational studies have found inverse associations between metformin use and survival rates of deadly cancers such as colon, liver, and lung cancers^23^. However, the results of epidemiological studies regarding use of metformin in patients with PCa have been inconsistent and provided suboptimal results^24^. Rothermundt *et al*.^25^ found stabilization and prolongation of prostate specific antigen (PSA) doubling time in 23 patients (52.3%) as well as an effect on metabolic parameters after starting metformin treatment. To date, however, the beneficial effect of metformin in reducing PCa incidence and improving overall survival is debated, particularly regarding the mode of action of metformin in clinical dosing in tumors. Furthermore, loss of LKB1 in T-cells results in the hyperactivation of the JAK/STAT signaling^26^, but there has been little evidence of a direct role of STAT3 downstream from metformin in t2DM PCa. In this study, we reveal an inverse correlation between STAT3 and mTORC1 expression in t2DM PCa patients undergoing metformin therapy. Using mouse genetics, we uncover for the first time the functional connections between STAT3 and LKB1/AMPK/mTORC1 signaling in PCa. We show that expression of LKB1/STK11 is directly regulated by STAT3 and regulates its transcriptional activity. Furthermore, we present results linking tumor proteomic profiling to the molecular mechanism of signal transduction to further dissect the effects of metformin on mTORC1 and CREB signaling in PCa. We also show that high CREB expression levels are strongly associated with a risk of biochemical and metastatic recurrence toward ADT-resistance in PCa patients. Finally, we link the STAT3 and CREB expression status to AR signaling and ADT-resistance in PCa patients, suggesting critical regulators and therapeutic targets of metastatic PCa.

## Results

Treatment of PCa with ADT leads to a metabolic syndrome that can contribute to cancer-related morbidity and mortality. Metformin can reverse the effects of the metabolic syndrome but also was shown to have a potential antineoplastic effect in several malignancies^27^. To investigate the clinical relevance of STAT3-mTORC1 signaling in human PCa, we performed antibody staining for STAT3 and the mTORC1 substrates p-4E-BP1, 4E-BP1 and p-S6 in tissue microarrays (TMA) of patients diagnosed with t2DM who underwent a radical prostatectomy due to organ confined PCa as described previously^28^. Patients were either treated with metformin (n=61) or insulin (data not shown), and as a control group we used patients who were not treated pharmacologically but only by diet procedures (n=41). Patient characteristics of the cohort were described previously^28^. In total, 570 tissue samples from 92 patients were collected. Embedded hepatic cells as well as prostate cell lines (LNCaP and PC3) were used as controls. PCa samples were prognostically scored using the Gleason score (GSC) with good prognosis less than or equal to a score of 7a and poor prognosis greater of equal to 7b. We observed higher STAT3 and lower p-4E-BP1 protein expression levels in the metformin-administered group GSC≥7b (n=20) compared to patient group GSC≥7b (n=26), which did not receive metformin (**Fig. 1A**). These data are consistent with the established role of metformin as an inhibitor of mTORC1 signaling through AMPK and tuberous sclerosis complex (TSC)^20^. The expression levels of other downstream substrates functionally controlling mTORC1 complex such as total 4E-BP1 and phospho-S6 levels remained unchanged, suggesting that alternative oncogenic events likely contribute to the signaling effects of metformin (**fig. S1A**).

**Fig. 1.**
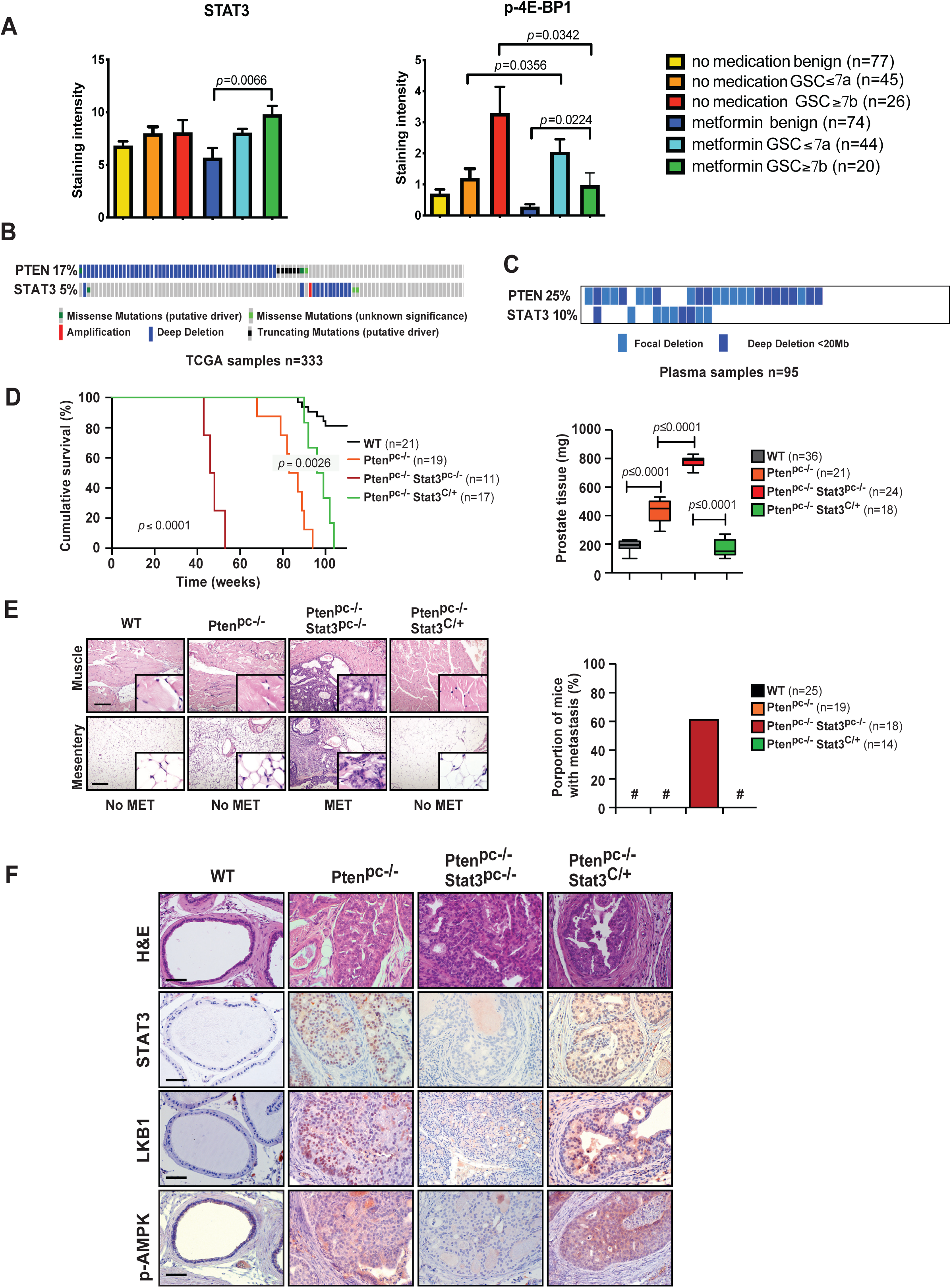
Loss of STAT3 accelerates metastatic progression and exhibits decreased LKB1/AMPK signaling. (**A**) IHC of radical prostatectomy specimens (benign and cancer core) stained with STAT3 and p-4E-BP1. TMA including benign and cancerous tissue with either metformin or diet =procedures as well as of patients without antidiabetic medication was employed (n=92) as described previously^28^. (**B**) Frequent homozygous and heterozygous deletions of *PTEN* and *STAT3* in primary PCa and mCRPC samples from TCGA database (n=333). (**C**) High occurrence of *PTEN* and *STAT3* deletions in plasma samples of PCa patients with aggressive PCa (n=95) (**D**) Kaplan–Meier cumulative survival analysis revealed a significant (*p*=0.0026; log-rank test) increase in lifespan of *Pten^pc−/−^Stat3^C/+^*compared to *Pten^pc−/−^Stat3^pc-/-^*mice; WT and *Pten^pc−/−^*mice served as controls (*n*=68). Prostate weights of 19-week- old WT, *Pten^pc−/−^*, *Pten^pc−/−^Stat3^pc−/−^*and *Pten^pc−/−^Stat3^C/+^*mice (n=99). Mean values are shown; Data were analysed by one-way analysis of variance with Tukey’s multiple comparison test; error bars: s.d. (**E**) IHC of muscle and mesentery, 52 weeks of age WT, *Pten^pc−/−^*, *Pten^pc−/−^Stat3^C/+^*and *Pten^pc−/−^Stat3^pc−/−^* mice. MET- metastasis, Scale bars, 100μm; insets: × 600 magnification. Percentage of mice with distant PCa metastases (*n*=76). (**F**) Haematoxilin/eosin (H&E) stains show only high- grade PIN in *Pten^pc−/−^Stat3^C/+^* mice compared with *Pten^pc−/−^* and *Pten^pc−/−^Stat3^pc−/−^*mice. Scale bars, 100μm. IHC analysis of Stat3, Lkb1 and p-Ampk in prostates from 19-week-old WT, *Pten^pc−/−^*, *Pten^pc−/−^ Stat3^pc−/−^*and *Pten^pc−/−^Stat3^C/+^* mice. Scale bars, 100μm.

### Loss of STAT3 accelerates metastatic progression and exhibits decreased LKB1/AMPK signaling

PTEN mutations are known to be important in the development of primary PCa. Interestingly, Oncomine mutation analysis revealed mutually exclusive deletions of PTEN and STAT3 in a large fraction of primary PCa (TCGA, n=333) (**Fig.1B**) in contrast to co-deletions in cfDNA plasma samples of patients with metastatic PCa (n=95)^29^, (**Fig.1C**, **fig. S1B**). The selective pressure on STAT3 loss during metastatic progression suggests that this step may be required to facilitate tumor dissemination. To test this hypothesis and identify pathways that trigger the metastatic program of PCa cells, we utilized a metastatic PCa mouse model with *Stat3* and *Pten* deletions (*Pten^pc−/−^ Stat3^pc-/-^*), which we previously described^8^. To broadly enable the role of STAT3 in metastatic reprogramming, we engineered constitutively active knock-in *Stat3C* allele replacing the endogenous WT allele^30^. The *Stat3^C/+^* mice were crossed with PCa mouse model (*Pten^pc−/−^)*^8^ animals to obtain *Pten^pc−/−^ Stat3^C/+^* mice. Constitutive activation of *Stat3* in *Pten^pc−/−^* mice, led to significant prolongation of survival and decreased prostate tissue weight (**Fig. 1D**) which is in sharp contrast to *Pten^pc−/−^ Stat3^pc−/−^* mice which died rapidly from metastatic disease. We could not find any evidence of tumor dissemination or metastasis in *Pten^pc−/−^ Stat3^C/+^*mice (**Fig. 1E**), up to >52 weeks of age, which suggests that functional Stat3 plays a major role in preventing metastatic dissemination. Furthermore, *Pten^pc−/−^ Stat3^C/+^* mice showed no signs of invasive tumor growth (**fig. S1C**) and only incidence of high-grade prostate intraepithelial neoplasia (PIN) (**Fig. 1F**), thereby impairing tumor progression and formation of metastasis. STAT3 has been described as a master metabolic regulator, which sustains glycolytic^10^ and oxidative phosphorylation^31^ activities of cells. The tumor suppressor LKB1, a key upstream regulator of metabolic sensor AMPK^32^, has also been shown to interact with the tumor suppressor PTEN^33^. Recently, Hermanova et al.^34^ intriguingly uncovered that the co-deletion of *Pten* and *LKB1* (*Stk11)* results in aggressive PCa tumors and formation of lung metastasis. This is reminiscent of the phenotype observed for deletions of *Pten and Stat3* in PCa mouse models. Importantly, we identified that *Pten^pc−/−^ Stat3^pc−/−^* tumors displayed reduced LKB1 and p-AMPK protein expression compared to *Pten^pc−/−^* tumors (**Fig. 1E**). In contrast, in *Pten^pc−/−^Stat3^C/+^* mice STAT3, LKB1 and p-AMPK overall protein levels as well as the number of positive staining cells were markedly increased in comparison to *Pten^pc−/−^ Stat3^pc−/−^* tumors (**Fig. 1E, fig. S1D**), suggesting that STAT3 is a regulator of LKB1/AMPK signaling.

### STAT3 and LKB1 cooperate to suppress mTORC1

To further corroborate the functional role of STAT3 in regulation of LKB1/AMPK/mTORC1, we analyzed passage-matched *Stat3^-/-^* MEFs. Loss of *Stat3* resulted in reduced LKB1/STK11 protein and mRNA levels, consistent with mTORC1 upregulation (**Fig. 2A**). Notably, knockdown of *STAT3* caused decreased LKB1/STK11 protein and mRNA expression in 22Rv1 PCa cells (**Fig. 2B and 2C**), indicating a possible STAT3-mediated transcriptional regulation of LKB1. Indeed, we confirmed a putative STAT3/consensus GAS binding site present in the LKB1 (STK11) promoter using the Transcription Factor Affinity Prediction Software (TRAP) and also validated by Linher-Melville et al.^35^. We performed a ChIP assay using an antibody against STAT3 in control versus STAT3 knockdown 22RV1 cells stimulated with IL-6. These results confirmed that endogenous STAT3 binds to the predicted regions on the LKB1 (STK11) promoter and that STK11 is a direct target gene of STAT3 (**Fig. 2D**). One of the major growth regulatory pathways controlled by LKB1–AMPK is the mTOR pathway. LKB1 activates AMPK which then rapidly inhibits a central integrator of cell metabolism and growth mTORC1^36^. mTORC1 is deregulated in most human cancers^37^ including genetic alterations of the mTOR signaling detected in 42% of primary and 100% of metastatic PCa^38^. Exploration of the functional relationship between STAT3 and mTORC1 signaling in the metastatic PCa mouse model revealed a significant increase in protein levels of mTORC1 downstream substrates p- 4E-BP1, 4E-BP1 and the phosphorylated ribosomal protein S6 in *Pten^pc−/−^ Stat3^pc−/−^* mice compared to *Pten^pc−/−^ Stat3^C/+^* mice (**Fig. 2E, F**), supporting a role for STAT3 in regulating mTORC1 signaling in PCa. In line with previous findings, we observed an increase of the *Ampk*α and the key members of *Lkb1* complex *Mo25* and *Strada* mRNA levels in *Pten^pc−/−^ Stat3^C/+^* mice (**fig. S2**). Having identified STAT3 as a transcriptional regulator of LKB1, we evaluated whether deletion of *Lkb1 (Stk11)* in *Pten^pc−/−^* mice would lead to deregulated STAT3 or mTORC1 signaling. Since the combination of *Pten* and *Lkb1* knock-out with heterozygous loss of *Pten* results in early lethality^34^, we analyzed the *Pten^pc+/−^ Stk11^-/-^* mouse model which had invasive PCa and extensive lung metastases with an incidence of >80%. Surprisingly, *Pten^pc+/−^ Lkb1^-/-^* tumors and lung metastases had elevated STAT3 and S6, as well as 4E-BP1 phosphorylation levels in comparison to *Pten^pc+/−^*, suggesting that tumors with a loss of LKB1/PTEN increase STAT3 expression/activity and mTORC1 activity, possibly through a negative feedback loop (**Fig. 2G, H**).

**Fig. 2.**
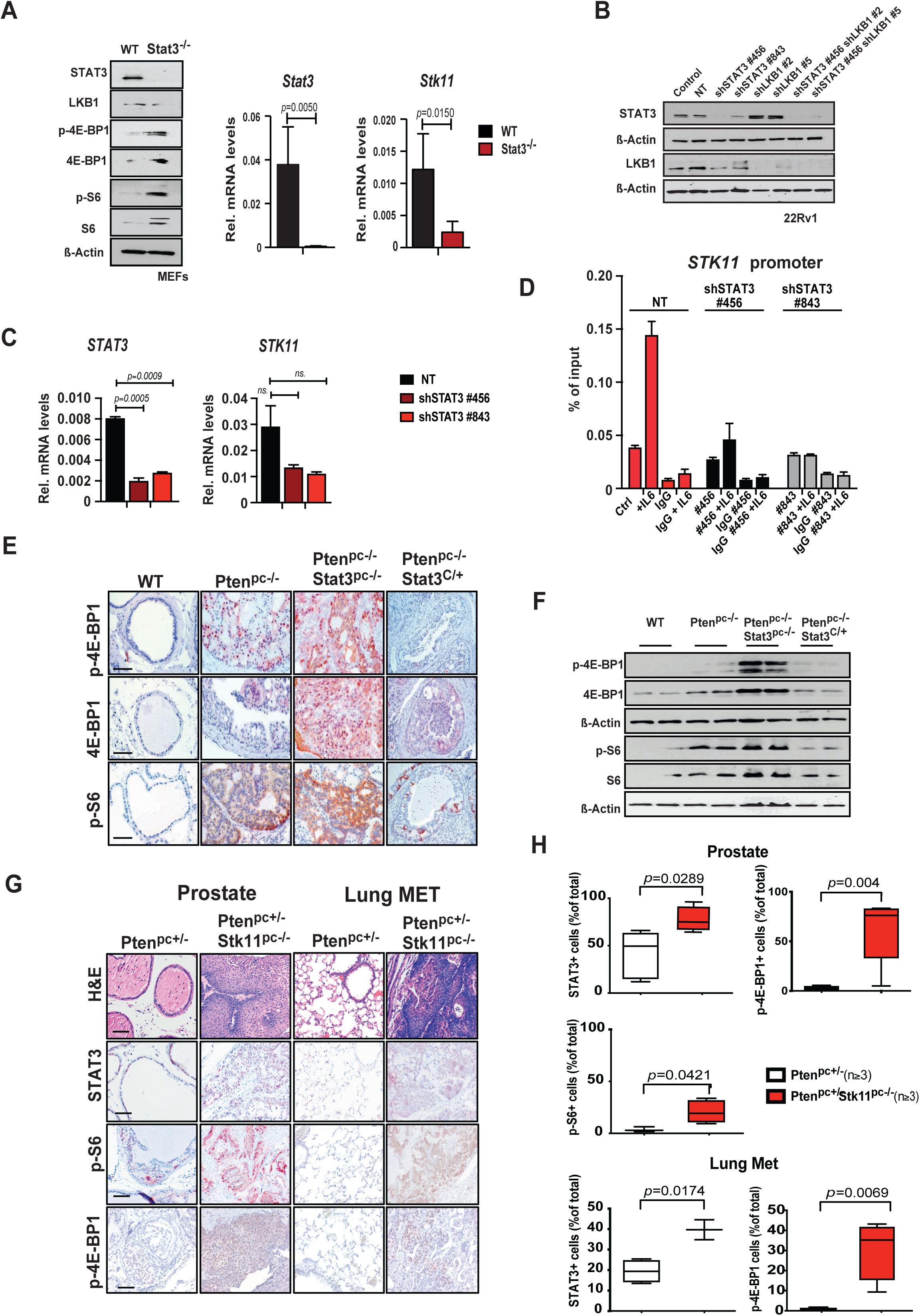
STAT3 and LKB1 cooperate to suppress mTORC1. (**A**) Western blot analysis of STAT3, LKB1, p-4E-BP1, 4E-BP1, p-S6 and S6 expression in WT and Stat3KO MEFs. β-actin serves as a loading control. qRT–PCR analysis of *STAT3* and *STK11* transcript levels in WT and Stat3KO MEFs (n=3 each). (**B**) Western blot analysis of STAT3 and LKB1 in 22Rv1 cells transfected with non-targeting (NT) or shRNAs specific for STAT3 and/or LKB1. (**C**) qRT–PCR analysis of *STAT3* and *STK11* in 22Rv1 cells transfected with NT or shRNA specific for STAT3 (n=3 each). (**D**) ChIP analysis of STAT3 binding to *LKB1 (STK11)* promoter. 22Rv1 cells harboring NT or two different shSTAT3 constructs were stimulated with IL-6 and immunoprecipated with STAT3 antibody or IgG as a negative control. Bars represent mean ± s.d. from 2 technical replicates. Precipitated DNA is presented as % of input. (**E**) IHC analysis of p-4E-BP1, 4E-BP1, p-S6 in prostates from 19-week-old WT, *Pten^pc−/−^*, *Pten^pc−/−^ Stat3^pc−/−^* and *Pten^pc−/−^Stat3^C/+^* mice. Scale bars, 100μm. (**F**) Western blots analysis of p- 4E-BP1, 4E-BP1, p-S6 and S6 expression in prostates from 19-week-old WT, *Pten^pc−/−^*, *Pten^pc−/−^ Stat3^pc−/−^* and *Pten^pc−/−^Stat3^C/+^*mice. β-actin serves as a loading control. (**G**) H&E and IHC analyses of Stat3, p-S6 and p-4E- BP1 in prostates and lung metastases from *Pten^pc+/−^*and *Pten^pc+/−^ Stk11^pc−/−^* mice, scale bars, 100 μm. (**H**) Quantification of cells positive for STAT3, p- 4E-BP1, and 4E-BP1 in 19-week-old *Pten^pc+/−^* and *Pten^pc+/−^ Stk11^pc−/−^*prostate tissue or lung metastases using HistoQuest software (n=3). Data were analysed by Student’s *t*-test and are shown as mean ± s.d.

### STAT3 is a central regulator of mTORC1 and CREB signaling

To identify the key oncogenic or tumor suppressor pathways involved in STAT3-dependent regulation of metastatic progression, we performed quantitative laser microdissection (LMD) and Label-free protein quantification (LFQ) proteome profiling on laser microdissected FFPE tumors of WT, *Pten^pc−/−^*, *Pten^pc−/−^ Stat3^pc−/−^* and *Pten^pc−/−^ Stat3^C/+^* prostate tissues. We detected 2,994 proteins that were altered between the *Stat3* deleted and constitutively active tumors. (**Fig. 3A**). We identified very low STAT3 protein expression in *Pten^pc−/−^Stat3^pc−/−^* compared to high expression in *Pten^pc−/−^Stat3^C/+^* prostate samples (**Fig. 3B**). It has been shown that STAT3 loss leads to disruption of mitochondrial metabolism and regulates the expression of genes involved in the mitochondrial oxidative phosphorylation (OXPHOS)^39^. Conversely, Pten haploinsufficiency results in mitochondrial dysfunction and increased activities of I-V mitochondrial complexes^11, 40^. We confirmed that *Pten^pc−/−^Stat3^pc−/−^* tumor cells show high-grade mitochondrial structural damage not evident in *Pten^pc−/−^Stat3^C/+^* prostate epithelial cells (**fig. S3A**). Similar inclusions have been described in defects in the assembly of the ATP synthase enzyme complex at the inner mitochondrial membrane, where inclusion bodies and loss of mitochondrial cristae occur^41^. Our earlier^9^ and current data (**fig. S3B**) showed increased levels of TCA/OXPHOS proteins in *Pten^pc−/−^Stat3^pc−/−^* tumor cells compared to *Pten^pc−/−^* tumors, suggesting that STAT3 is a key component of dynamic mitochondrial bioenergetics and redox regulation that enables cells to maintain homeostasis and energy metabolism under tumorigenesis and metastatic energetic stress. To explore differentially expressed tumor intrinsic signaling pathways in advanced PCa, we analyzed STAT3-dependent signaling profile at the level of proteome. We observed that TIMP metallopeptidase inhibitor 1 (TIMP1), Vimentin (VIM) and fascin actin-bundling protein 1 (FSCN1) were over- expressed in the context of *Stat3* activation. TIMP1 is a known STAT3 downstream target gene^42^ and other studies suggested that VIM^43^ and FSCN1^44^ are regulated through STAT3 phosphorylation and could be possible STAT3 targets. Transcriptional activation by STAT3 proteins has been shown to require the recruitment of CREB-binding protein (CBP)/p300 coactivators. In line with these results, CBP/p300 can interact with the activation domain of STAT3 to regulate transcription^45^. A recent study by Li et al.^46^, demonstrated that CREB activity is required for mTORC1 signaling in primordial follicles. This coexisting activation of CREB and mTORC1 activity remains unclear in cancer. Here, we performed comprehensive global protein expression profiling which showed an enrichment of mTOR and CREB signaling pathways (**Fig. 3C**) in *Pten^pc−/−^Stat3^pc−/−^* tumor cells, suggesting a critical role of STAT3 for mTORC1 and CREB signaling in tumorigenesis and metastatic formation. To assess the effect of STAT3 inhibition on mTOR and CREB signaling *in vivo*, we treated human LNCaP xenografts with JAK/STAT inhibitor Ruxolitinib. As for *Pten^pc−/−^ Stat3^pc-/-^* PCa tumors we observed significant upregulation of p-CREB, CREB, p-4E-BP1, 4E-BP, p-S6 and S6 protein expression (**Fig. 3D**), suggesting that blockade of STAT3 signaling can positively affect CREB expression and phosphorylation along with mTORC1 signaling. Furthermore, western blot analysis revealed that knockdown of STAT3 increased p-CREB and CREB protein levels in 22Rv1 cells. Accordingly, STAT3 add-back in PC3 cells lacking STAT3 led to reduced p-CREB and CREB protein levels (**Fig. 3E**).

**Fig. 3.**
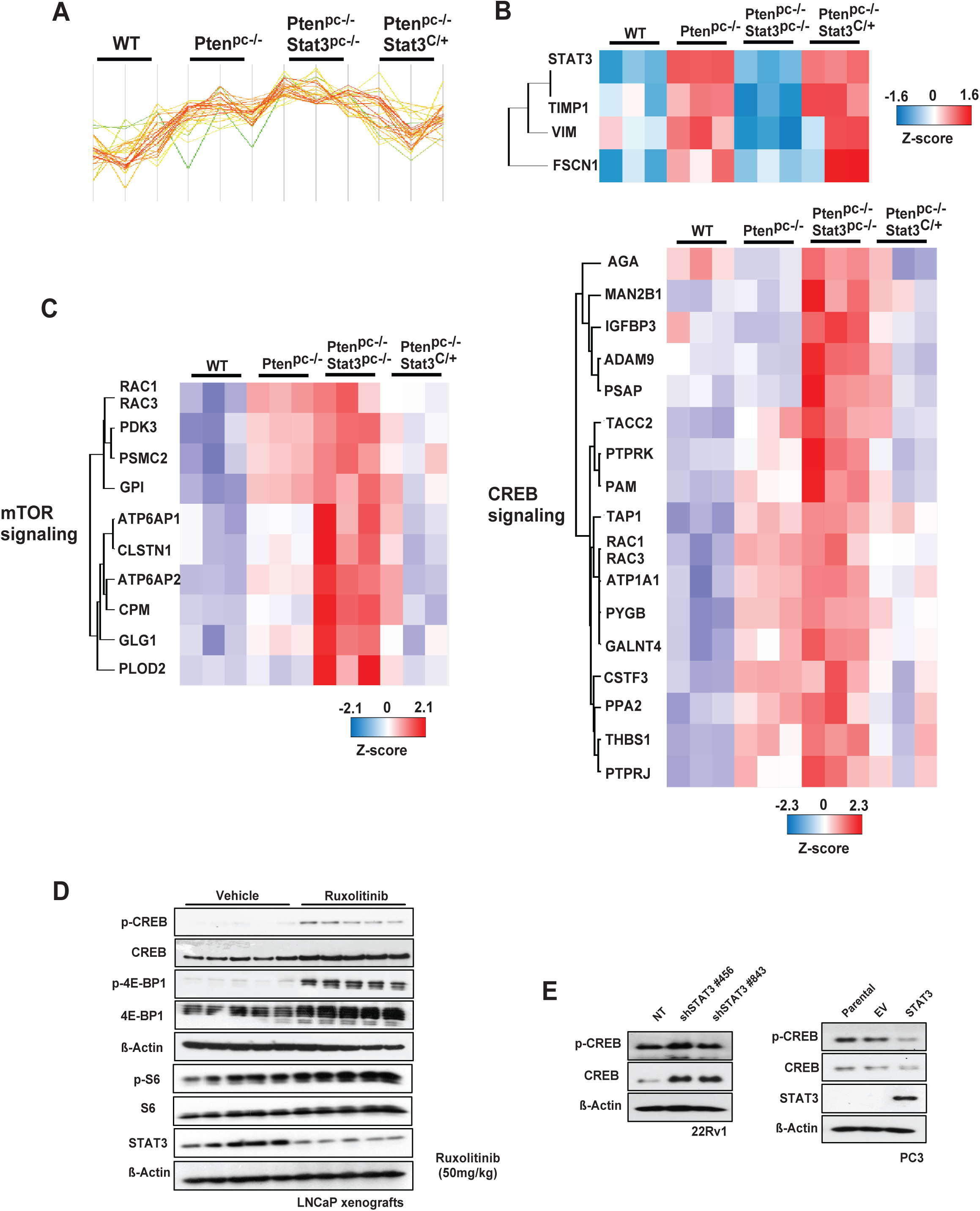
STAT3 is a central regulator of mTORC1 and CREB signaling. (**A, B**) Signature and heatmap of Stat3 and Stat3-regulated proteins in FFPE laser-microdissected prostates of 19-week-old WT, *Pten^pc−/−^*, *Pten^pc−/−^Stat3^pc−/−^*and *Pten^pc−/−^Stat3^C/+^* (n=3 each) using unbiased LC- MS/MS proteomics. (**C**) The heatmap shows reprogramming of mTOR and CREB metabolic pathways in PCa with significant enrichment (hypergeometric test, *q*-value□<□0.05). (**D**) Western blot analysis of p- CREB, CREB, p-4E-BP1, 4E-BP1, STAT3, p-S6 and S6 of LNCaP xenograft tumors treated with vehicle or Ruxolitinib (50 mg kg^-1^) for 22 days. β-actin serves as a loading control. (**E**) Western blot analysis of p-CREB and CREB in 22Rv1 cells transfected with non-targeting (NT) or shRNAs specific for STAT3. Western blot analysis of STA3, p-CREB and CREB in PC3 cells transfected with an empty vector (EV) or STAT3 add-back. PC3 cells lacks STAT3 expression.

### A clinically relevant dose of metformin inhibits CREB/mTORC1 in a manner that requires AR and STAT3 signaling

The frontline and most prescribed antidiabetic drug metformin also showed AMPK-dependent mTORC1 inhibition via TSC/RHEB and has been considered as a potential anticancer agent^47^. Metformin is still being evaluated as an adjuvant therapy in various disease settings^24^. To investigate whether the STAT3 status could directly affect metformin sensitivity through modulation of CREB and mTORC1 signaling, we analyzed the antiproliferative effects of metformin. We found that metformin treatment in a clinically relevant dose^48^ decreased size, weight, and tumor volume in AR/STAT3 expressing 22Rv1 cells but was ineffective for PC3 xenografts *in vivo* (**Fig. 4A-C**). Metformin also significantly reduced cell viability in all PCa cell lines except the PC3 cell line, which is androgen insensitive and is known to lack AR and STAT3^49^ (**Fig. 4D**). Metformin inhibits mitochondrial electron transport chain (ETC) complex I as well as other targets of mitochondrial metabolism^50^. Furthermore, it has been shown that acute and chronic low dose metformin treatment effectively impacted the cytosolic/mitochondrial redox state and inhibited mitochondrial glycerol phosphate dehydrogenase (mGPD)^51^. Since both complex I inhibition and mGPD decreased activity could lead to NAD^+^ deficiency, we measured NAD^+^, NADH and ATP in 22Rv1 and PC3 cells *in vivo* using HRSM and observed decreased levels of NAD^+^ in PC3 cells (**fig. S4A**) consistent with an altered NAD+ regeneration and sensitivity leading to malignant phenotypes by promoting clonal cell growth and migration upon loss of STAT3 in triple negative breast cancer^52^. In contrast to 22Rv1, PC3 tumors treated with metformin exhibited high proliferative counts of Ki-67^+^ cells associated with an increased number of CC3^+^ apoptotic cells (**fig. S4B**). Importantly, we observed that metformin treatment suppressed mTORC1 and CREB signaling in STAT3-expressing 22Rv1 xenografts while the ability of metformin to suppress p-4E-BP1, 4E-BP1, p-S6, S6, p-CREB and CREB was greatly diminished in PC3 STAT3-deficient tumors (**Fig. 4E)**, underlying the mTORC1 and CREB dependency on STAT3 status. Similarly, induction of LKB1 was observed in STAT3-expressing 22Rv1 xenografts treated with metformin while blunted in PC3 STAT3-deficient tumors. We next determined whether STAT3 is critical for sustained AR signaling promoted by LKB1/AMPK and CREB activation. Inhibition of STAT3 in LNCaP xenografts treated with Ruxolitinib led to induction of AR and PSA levels (**Fig. 4F**), potentially indicating emergence, and increasing prevalence to ADT-resistance. On the other hand, metformin treatment repressed AR and PSA levels in 22Rv1 xenografts while PC3 tumors lacked AR and PSA protein expression (**Fig. 4G**). These results indicate that STAT3 and CREB signaling are critical for AR regulation and potential ADT-resistance.

**Fig. 4.**
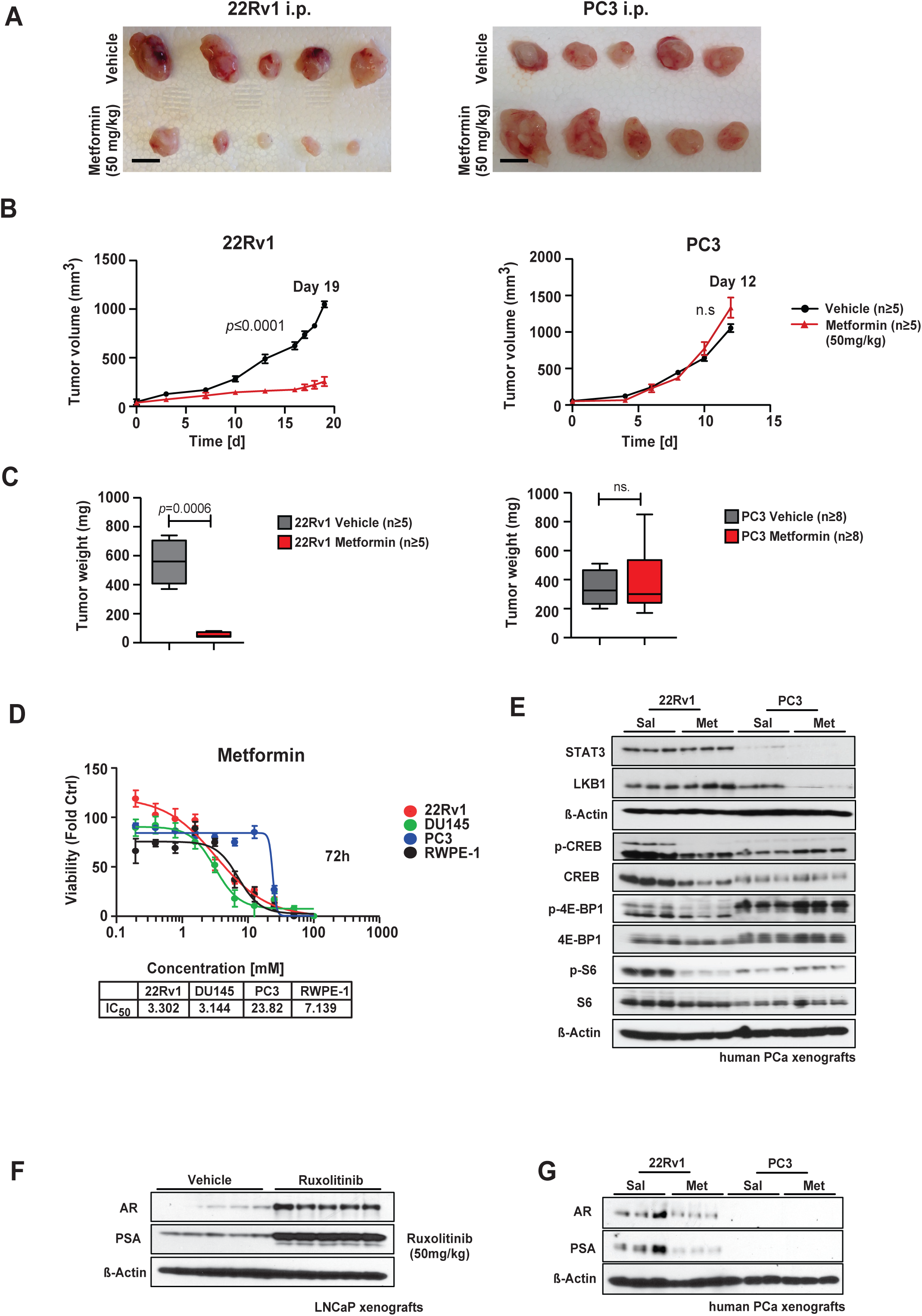
Clinically relevant dose of metformin inhibits CREB/mTORC1 in a manner that requires AR and STAT3 signaling. (**A)** Gross anatomy assessment of representative 22Rv1 and PC3 xenograft tumors i.p. treated with vehicle or metformin (50mg/kg). Scale bars, 10 mm. Mean values are shown; error bars: s.d. (n=5). (**B**) 22Rv1 and PC3 cells were implanted subcutaneously in mice and grown until tumors reached the size of approximately 100 mm^3^. Xenografted mice were randomized and then received (n=5 per group) vehicle or 50mg/kg metformin i.p. daily. Mean tumor volume ± s.d. is shown. (**C)** Tumor weights assessment of representative 22Rv1 and PC3 xenograft tumors i.p. treated with vehicle or metformin (50mg/kg). Scale bars, 10 mm. Mean values are shown; error bars: s.d. (n=5). (**D**) Comparison of IC_50_ values of metformin for human PCa cell lines (22Rv1, DU-145 and PC3) and untransformed human prostate cell line RWPE-1. **E)** Western blot analysis of STAT3, LKB1, p-CREB, CREB, p-4E- BP1, 4E-BP1, p-S6 and S6 in 22Rv1 and PC3 xenograft tumors. β-actin serves as a loading control. (**F**) Western blot analysis of AR and PSA in LNCaP xenograft tumors treated with vehicle or Ruxolitinib (50 mg/kg) for 22 days. β-actin serves as a loading control. Western blot analysis of AR and PSA in 22Rv1 and PC3 xenograft tumors i.p. treated with vehicle or metformin (50mg/kg). β-actin serves as a loading control.

Moreover, *Stat3^-/-^* MEFs (**fig. S5A**) and *Pten^pc−/−^ Stat3^pc−/−^* mice (**fig. S5B**) showed elevated p-CREB and CREB protein expression compared to control. Taken together, these results demonstrate that in PCa mTORC1 and CREB signaling are regulated by LKB1-AMPK in an STAT3-dependent manner and may control PCa cell growth and metastatic spread.

### STAT3 and PTEN are negatively correlated with mTORC1 in PCa

Because loss of STAT3 and PTEN in our model systems demonstrated upregulation of mTORC1 relative to primary and metastatic PCa, we explored publicly available data sets of prostate carcinomas and metastatic PCa from Arredouani et al.^53^ and Lapointe et al.^54^. The cohorts revealed significant mRNA downregulation of *STAT3*, *PTEN* and a downstream target of LKB1 the salt-inducible kinase 3 (SIK3) expression and upregulation of *EIF4EBP1* (4E- BP1) (**Fig. 5A-D**). Similar results confirmed clinical failure and more likely development of distant metastasis occurring in patients^55^ in the presence of low PTEN and high EIF4EBP1 and EIF4E mRNA expression (**fig. S6A**). Focusing on the SIK3 mRNA expression in PCa, we also analyzed a previously published PCa dataset^38^. PCa patients with low SIK3 expression were significantly more likely to have a lower probability of progression-free survival (**fig. S6A**). We focused our analysis on TMA of PCa patient cohorts (n=83). In line with our findings in mouse model systems and patient data sets, low STAT3 and high 4E-BP1 and p-4E-BP1 cytoplasmic and nuclear protein expression were seen in prostate carcinomas (**Fig. 5E**, **fig. S6B**). Further research shows that increased p-4E-BP1 protein expression is present in advanced Gleason 5 carcinoma compared to Gleason 3 or 4 (**fig. S6B**). In summary, our findings from PCa patient samples mimic the molecular phenotype of preclinical mouse models and engineered cell lines and exhibited reprogramming of LKB1/SIK3 and mTORC1 metabolic pathways.

**Fig. 5.**
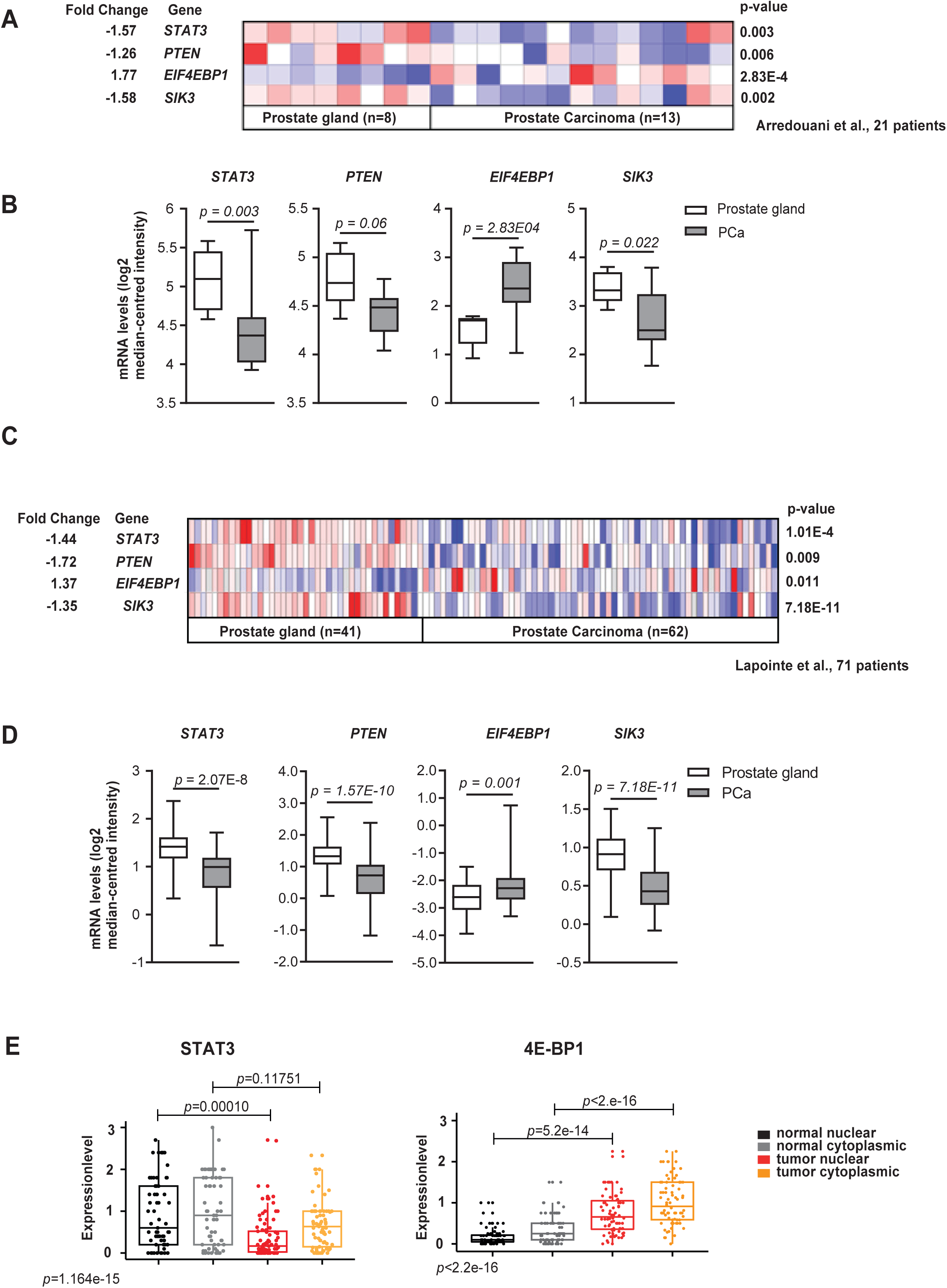
STAT3 and PTEN are negatively correlated with mTORC1 in PCa. (**A**) Heatmaps depicting significant downregulation of *STAT3, PTEN* and *SIK3* mRNA levels and concomitant upregulation of *EIF4EBP1* mRNA expression in prostate carcinoma patient samples (n=13) compared with healthy prostate gland tissues (n=8). Colors were normalized to depict relative mRNA expression values (log2 median-centered intensity) within each row; dark blue represents the lowest relative expression level and dark red represents the highest relative expression level. Data were extracted from the Oncomine™ Platform (*78*) and from the Arredouani Prostate study (*53*). (**B**) Gene expression levels depicting significant downregulation of *STAT3* (-1.57-fold)*, PTEN* (-1.26-fold) and *SIK3* (-1.58-fold) mRNA and concomitant upregulation of *EIF4EBP1* mRNA (1.77-fold) in prostate carcinoma patients (n=13) compared with normal prostate gland samples (n=8). Data (log2 median- centered intensity) were extracted from the Oncomine™ Platform from the Arredouani Prostate dataset. Representation: boxes as interquartile range, horizontal line as the mean, whiskers as lower and upper limits. (**C**) Heatmap depicting significant downregulation of *STAT3, PTEN* and *SIK3* mRNA levels and concomitant upregulation of *EIF4EBP1* mRNA expression in prostate carcinoma patients compared with normal prostate gland samples (log2 median-centered intensity). Data were extracted from the Oncomine™ Platform from the Lapointe Prostate dataset. (**D**) Gene expression levels depicting significant downregulation of *STAT3* (-1.44-fold)*, PTEN* (-1.72-fold) and *SIK3* (-1.35-fold) mRNA and concomitant upregulation of *EIF4EBP1* mRNA (1.37-fold) in prostate carcinoma patients (n=59-62) compared with normal prostate gland samples (n=37-41). Data (log2 median-centered intensity) were extracted from the Oncomine™ Platform from the Lapointe Prostate dataset. Representation: boxes as interquartile range, horizontal line as the mean, whiskers as lower and upper limits. (**E**) Boxplots representing protein expression of STAT3 and 4E-BP1 in cytoplasmatic or nuclear stainings detected by IHC in normal-like glands or tumors in PCa patient TMAs (n=83).

### CREB signaling predicts ADT-resistance and metastatic progression in PCa patients

Considering these findings, we determined the clinical relevance of CREB signaling in patients affected by PCa by analyzing TMA of 83 patient cases. IHC analysis revealed that the majority of tumors (**Fig. 6A)**, and PCa patients with GSC 8-10 expressed high protein levels of CREB (**fig. S7A**). Next, we determined whether CREB expression could predict worse clinical outcome. Indeed, we found that high CREB expression significantly correlates with inferior BCR-free survival in PCa patients (**Fig. 6B, fig. S7A**). On multivariable analysis, a high CREB1 gene expression pattern was associated with an increased risk of biochemical recurrence (HR = 1.56 [1.13–2.16]; *p* = 0.0703; (**Fig. 6C)** and metastatic recurrence (HR = 2.09 [1.15–3.83]; *p* = 0.0164; (**Fig. 6D)**. Since CREB1 mRNA expression predicts metastatic recurrence better than biochemical recurrence this suggests that metastatic subgroup of PCa patients with high CREB expression have a greater risk of developing metastatic disease progression. The SIK3 expression in tumor cells and PCa patients remain unchanged (**fig. S7, C**). Taken together, these data suggest that targeting CREB signaling may be the optimal approach for suppressing metastatic PCa in the molecular context of loss of *PTEN*. Next, we sought to compare transition from androgen-dependent (AD) to an androgen-independent (AI) PCa in LNCaP xenograft models during serial propagation in castrated mice^56^ (**Fig. 6E)**. We observed a clear upregulation of AR and p- CREB protein levels and STAT3 downregulation in AR-independent LNCaP xenografts linking the CREB expression to distinct tumorigenic behavior of CRPC and suggesting therapeutic regiments targeting CREB in mCRPC.

**Fig. 6.**
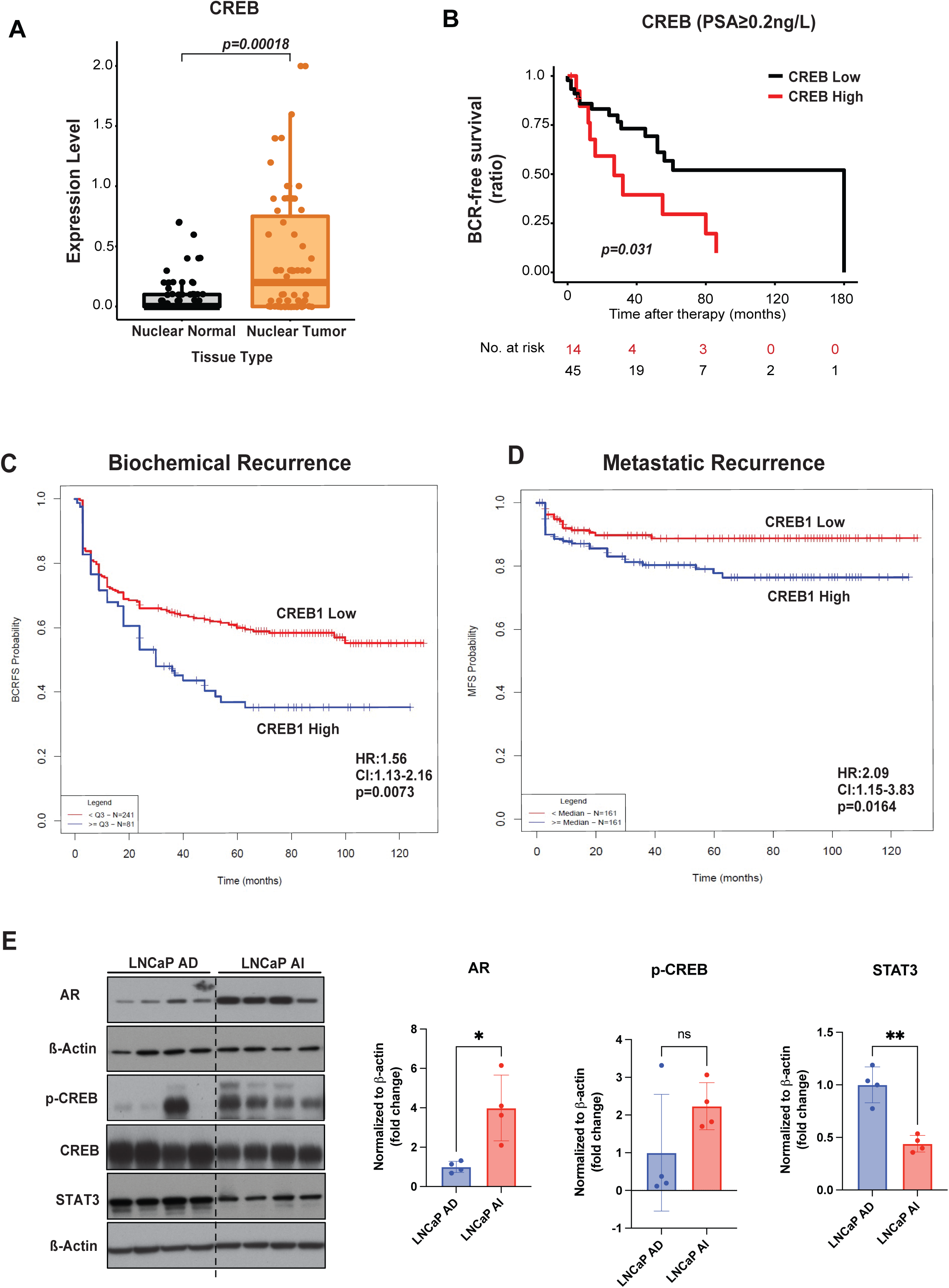
CREB signaling predicts ADT-resistance and metastatic progression in PCa patients. (**A**) Boxplots representing protein expression of CREB in nuclear stainings detected by IHC in normal-like glands or tumor cells in PCa patient TMAs (n=83). (**B**) Kaplan–Meier analysis of BCR-free survival ratio based on CREB protein expression in a panel of 83 PCa patients (PSA > 0.2ng/L). (**C**) Association of CREB expression at predicting time to biochemical recurrence of high/low-risk disease in the resection cohort. Reduced progression-free survival in months of the “high-risk” subgroup (blue) of 112 patients when compared with the “low-risk” subgroup (red) of 125 patients (HR = 1.56 [1.13–2.16]; *p* < 0.0073). (**D**) Association of CREB expression at predicting time to metastatic disease recurrence of high/low-risk disease in the resection cohort. Reduced progression-free survival in months of the “high-risk” subgroup (red) of 112 patients compared with the “low-risk” subgroup (blue) of 125 patients (HR = 2.09 [1.15–3.83]; *p* < 0.0164). HR = hazard ratio. (**E**) LNCaP xenograft model was serially passaged in castrated NSG males. WB analysis of AR, p-CREB, CREB, STAT3 protein expression in LNCAP AD/AI tumors. β-actin serves as a loading control.

## Discussion

The initiation and progression of PCa is driven by AR signaling. ADT constitutes a pillar of systemic therapy for patients with advanced PCa. However, over the past years several studies have shown that up to 60% of patients with advanced PCa have molecular alterations in non-AR related pathways^57^. In particular, mutations in the genes encoding components of the DNA damage response and repair such as *BRCA1* and *BRCA2*, are present in a significant proportion of patients with PCa^58^. Poly(adenosine diphosphate–ribose) polymerase inhibitors (PARPi) are the first approved targeted therapy for metastatic castration-resistant prostate cancer (mCRPC) with previous second-generation ADT in patients with BRCA1/2 mutations (Europe) or ATM alterations (USA). Although there has been interest to further exploit other genetic alterations therapeutically, the molecular machinery and cellular energetic elements governing disease progression remains poorly characterized. Tumor histology, PSA and imaging still represent key factors in adjuvant therapy decisions, but suffer from poor reproducibility, particularly among non-subspecialist pathologists. Currently, there is no infallible method of distinguishing aggressive from indolent tumors. However, breakthrough discoveries during the past century have profoundly altered the clinical management for patients with PCa. Classification of disease subgroups based on computational histological pattern recognition and prediction of genomic features is now available for PCa prognostication^59^. Metabolic syndrome is characterized by insulin resistance, hyperinsulinemia, hyperlipidemia, and obesity and is associated with increased development and progression of aggressive PCa^24^. The metabolic syndrome is present in more than 50% of the men undergoing long-term ADT (at least 12 months) and results not only in insulin resistance, but also in hyperglycemia^60^. Therefore, it became an important factor for biochemical failure after prostatectomy and radiotherapy. In a retrospective study, Flanagan *et al*.^61^ reported that metabolic syndrome was associated with a shorter time to PSA progression and inferior overall survival in patients with PCa receiving ADT. Metformin is still the glucose-lowering drug of choice for the management of t2DM. Metformin restores insulin and glycemic metabolic balance and according to preclinical studies *in vitro* and *in vivo* inhibits growth of cancer cells^62^. In a recent study, Gutkind *et al*.^63^ showed in a phase II trial of metformin in individuals with oral premalignant lesions that metformin administration results in inhibition of proliferation and decreased mTOR activity (pS6 IHC staining). Similarly, we also observed low p-4E-BP1 protein expression levels in the metformin-administered group GSC≥7b compared to PCa patient group GSC ≥ 7b who did not receive an antidiabetic drug.

However, metformin administration in PCa patients led to steadily increasing levels of STAT3 expression in the metformin-administered group suggesting that a long-term metformin treatment may regulate multiple biological functions and signaling during malignant transformation. In this scenario, in analogy to IL-6 and its downstream STAT3 in regulating senescence-inducing circuit controlled by p53-ARF^8^ and secretion of proinflammatory cytokines and chemokines^64^, cancer cells treated with metformin rewire the metabolic and signaling pathways in response to mTOR and PI3K/AKT inhibition. Importantly, our data revealed that a long-term effect of metformin administration requires STAT3 for mTORC1-mediated inhibition, indicating that JAK/STAT signaling represents the primary signaling mechanism involved. Despite a well-accepted oncogenic role, STAT3 can also exert tumor suppressor activity in the context of tissue type^7, 8^. Previous evidence demonstrates that the tumor-suppressive function of STAT3 is intimately linked to PTEN function and PI3K/AKT activation in astrocytes^65^. In line with this finding, we found 5% of STAT3 and 17% PTEN deletions in PCa patients using the TCGA database, but a large number of concomitant deep deletions of PTEN and STAT3 in cfDNA plasma samples of patients with metastatic PCa suggesting that STAT3 loss is frequently associated with PTEN loss in advanced and metastatic PCa. Therefore, a functional relationship underlying synthetic interactions between STAT3 and PTEN might have an obligate role in effecting carcinogenesis and tumor maintenance. Using genetically engineered mouse models (GEMMs), we demonstrate that genetic inactivation of Stat3 in *Pten^pc−/−^* mice decreases the survival, while constitutive Stat3 activation caused tumor regression and extended survival beyond PTEN-deficiency. Of note, we observed metastatic formation in *Pten^pc−/−^*, *Stat3^pc−/−^* mice to similar extent as in *Pten^pc−/−^*, *Tp53^pc−/−^* or *Pten^pc−/−^*, *Smad4^pc−/−^*GEMMs^66, 67^ suggesting the importance of STAT3 signaling for metastatic risk in human PCa. LKB1/STK11 functions as a master regulator of cell metabolism and energy stress responses. LKB1 signaling negatively regulates tumor growth and metastatic formation in mouse models of lung cancer and melanoma^68^. Furthermore, we observed deregulation of LKB1/AMPK and mTORC1 signaling tightly associated with STAT3 status, indicating that STAT3 signaling is a critical regulator of PCa metastasis by switching senescence and control of LKB1-dependent biochemical signatures of metastatic behavior. In addition to the functional inactivation of LKB1/AMPK in *Pten^pc−/−^* tumors from the loss of *Stat3*, we used ChIP and confirmed that STAT3 is binding to the STK11 (LKB1) promoter. We found that *Stat3* knockout leads to upregulation of mTORC1 signaling. Of note, unbiased LMD and LFQ shotgun proteome profiling revealed a prominent mTOR and CREB signaling dependency in loss of *Stat3* tumors, which displayed metastatic phenotypes, suggesting a dominant regulatory function for mTOR and CREB in PCa metastasis formation. Consequently, treatment with Ruxolitinib in LNCaP xenograft tumors led to upregulation of CREB and mTORC1 signaling, thereby potentially promoting metastatic behavior. CREB forms a complex with CREB-binding protein (CBP) and recruits the transcription machinery at the gene promoter to initiate CREB-dependent gene transcription^69^. Given the evidence that metformin can stimulate CBP phosphorylation in a mouse model of insulin resistance^70^, we treated mice bearing STAT3-proficient and deficient PCa tumors with metformin. We observed CREB/mTORC1-dependent effects of metformin in STAT3- proficient tumors versus independent mechanisms seen in STAT3-deficient tumors. Zhang *et al*., showed that in aggressive neuroendocrine prostate cancers ADT promotes angiogenesis and neuroendocrine differentiation through the CREB-EZH2-TSP1 pathway^71^, which suggests that CREB may be a valuable target in mCRPC. This study suggests that certain genetic alterations may provide robust metastatic advantages and potentially rewire distinct metabolic pathways in metformin-treated cancer patients even from early stages of tumorigenesis, alleviating the need for additional cancer therapies to suppress metastatic potential. Of high translational impact we show here that increased CREB levels are frequently associated with tumor recurrence and metastatic disease in PCa patients and that patients with tumor expressing high CREB levels have shorter PSA relapse. Given the *in vivo* data in GEMMs and the clinical evidence of CREB expression in PCa patients, we believe that metformin should be administered with caution in patients with aggressive prostate cancers. It has been shown that metformin represses prostate cancer cell viability and enhances apoptosis by targeting the AR signaling pathway^72^. Moreover, we found that STAT3 is required for the inhibition of AR and PSA by metformin, which indicate STAT3 and CREB as a signaling node for ADT-resistance in metastatic PCa. Coupled with the original observation of increased STAT3 expression in metformin treated PCa patients, this expands the contexts in which metformin is traditionally thought of playing only antitumor activity. By contrast, upon ADT, these tumor cells upregulate CREB and mTORC1 signaling, which facilitates mCRPC progression. We also show that Ruxolitinib treatment enhanced AR and PSA protein levels. Since the majority of patients relapse with typical AR-positive adenocarcinoma with rising PSA levels, we now hypothesize that common JAK inhibitors for treatment of rheumatoid arthritis and polycythemia vera can potentially increase the risk of resistance to AR inhibitors in PCa. Increased PSA levels indicate the recurrence of CRPC, and the patient is then put on second-generation ADT (enzalutamide or abiraterone). Despite initial success, patients will eventually experience enzalutamide resistant secondary CRPC with metastatic lesions. Recently, Kim *et al*.^73^ showed that pharmacological inhibition of CREB1 with the specific CREB inhibitor 666-15 abrogated the number of formed lung metastases and reduced pancreatic ductal adenocarcinoma metastasis. Collectively, our results suggest a proof-of-concept study to access objective response and efficacy targeted CREB inhibition in mCRPC resistant to second generation ADT.

Previous evidences demonstrate that STAT3 plays a pivotal role in tumor-infiltrating immune cells^74^ and regulates the levels of several secreted factors such as interferons, cytokines and growth factors^75^, thereby exerting profound immune effects. In a recent report, Pore *et al*., showed that reduction of STAT3 in the tumor microenvironment using an anti-sense oligonucleotide reversed immunotherapy resistance in preclinical STK11 knockout models^76^. Several developments in the past have advanced CRPC treatment, including enzalutamide and abiraterone, however mCRPC has been very resistant to checkpoint inhibitors, because it is immunologically cold with a few tumor-infiltrating T cells. Given the role of CREB signaling in regulation of several immune-related genes including IL-2, IL-6, IL-10 and TNF^77^, it is tempting to speculate that CREB signaling downstream of these effectors in immune cells mediates immunosuppressive effects and resistance to immunotherapy in CRPC patients. Together, our findings reveal insights into how STAT3 controls metastatic PCa formation and support the concept that CREB is a potential target for interventions in patients with mCRPC.

---

**Supplementary Fig. 1.**
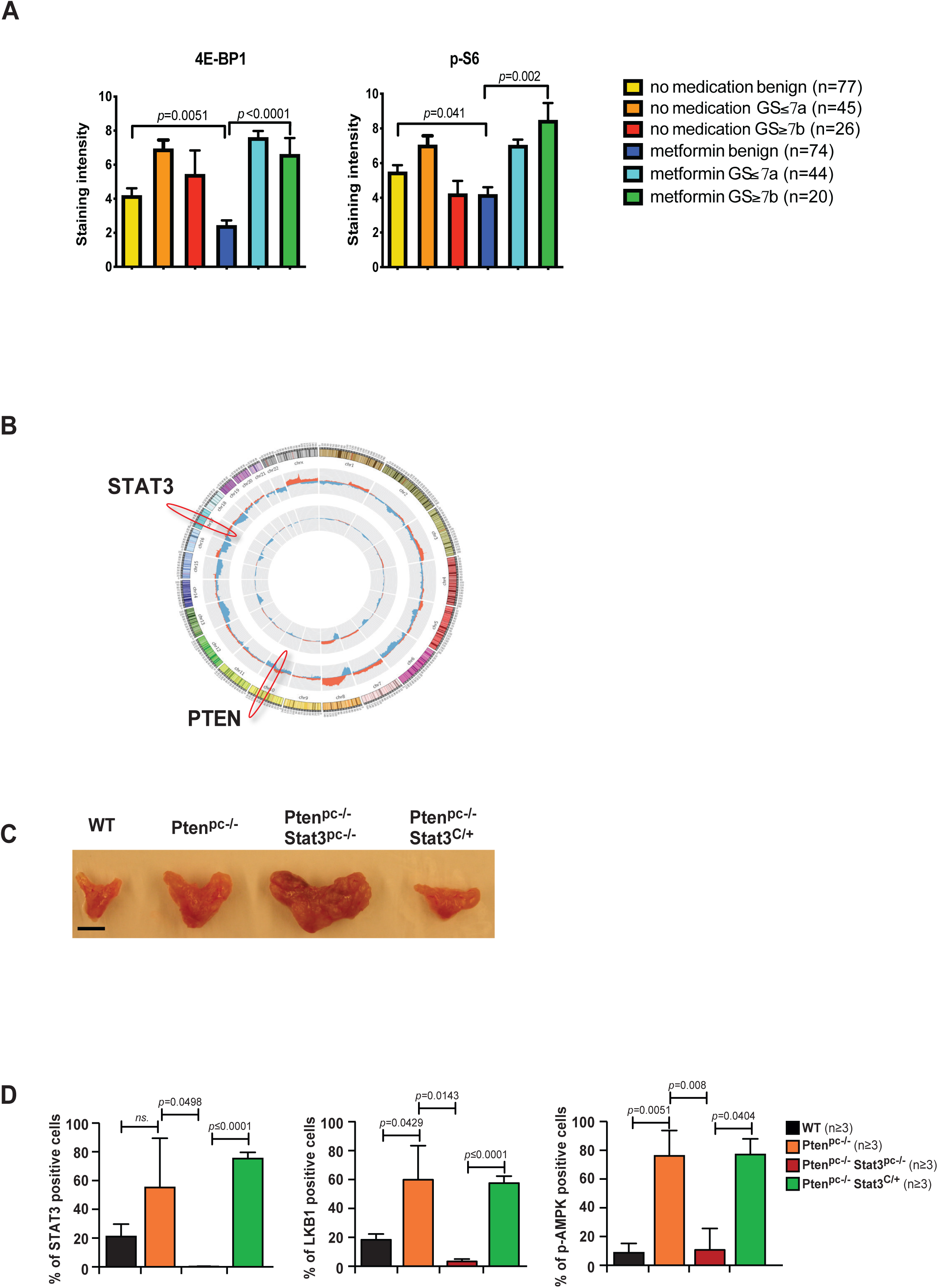
(**A**) IHC analysis of radical prostatectomy specimens (benign and cancer core) of 118 PCa patients stained with 4E-BP1 and p-S6. 61 used metformin and 47 PCa patients with no pharmacological treatment were included as a control group. (**B**) Circos plot of STAT3 and PTEN deletions: inner circle TCGA (n=492) and outer circle plasma-DNA samples from advanced PCa patients (n=43). (**C**) Gross anatomy of representative prostates isolated at 19 weeks of age from WT, *Pten^pc−/−^*, *Pten^pc−/−^Stat3^pc−/−^*and *Pten^pc−/−^Stat3^C/+^* mice. Scale bars, 10□mm. (**D**) Bar graphs indicate percentage of cells positive for Stat3, Lkb1 and p-Ampk in prostates of 19-week-old WT, *Pten^pc−/−^*, *Pten^pc−/−^Stat3^pc−/−^*and *Pten^pc−/−^Stat3^C/+^*. Protein levels quantification was done with HistoQuest software (n=3).

**Supplementary Fig. 2.**
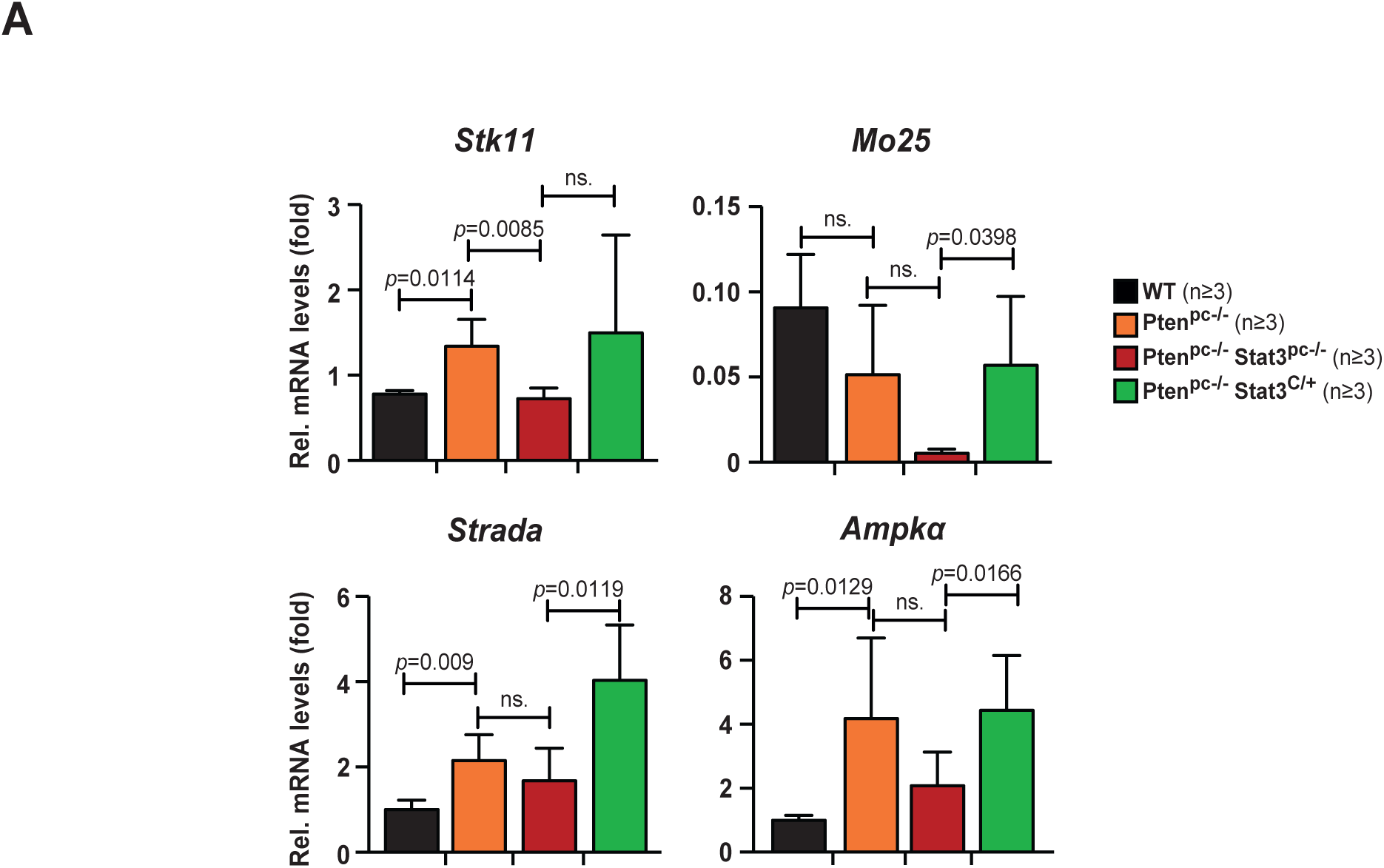
qRT–PCR analysis of *Stk11, Mo25, Strada and Ampk*α mRNA expression in prostates of 19-week-old WT, *Pten^pc−/−^*, *Pten^pc−/−^Stat3^pc−/−^* and *Pten^pc−/−^Stat3^C/+^*(n=3 each). Data were analyzed by Student’s *t*-test and are shown as mean ± s.d.

**Supplementary Fig. 3.**
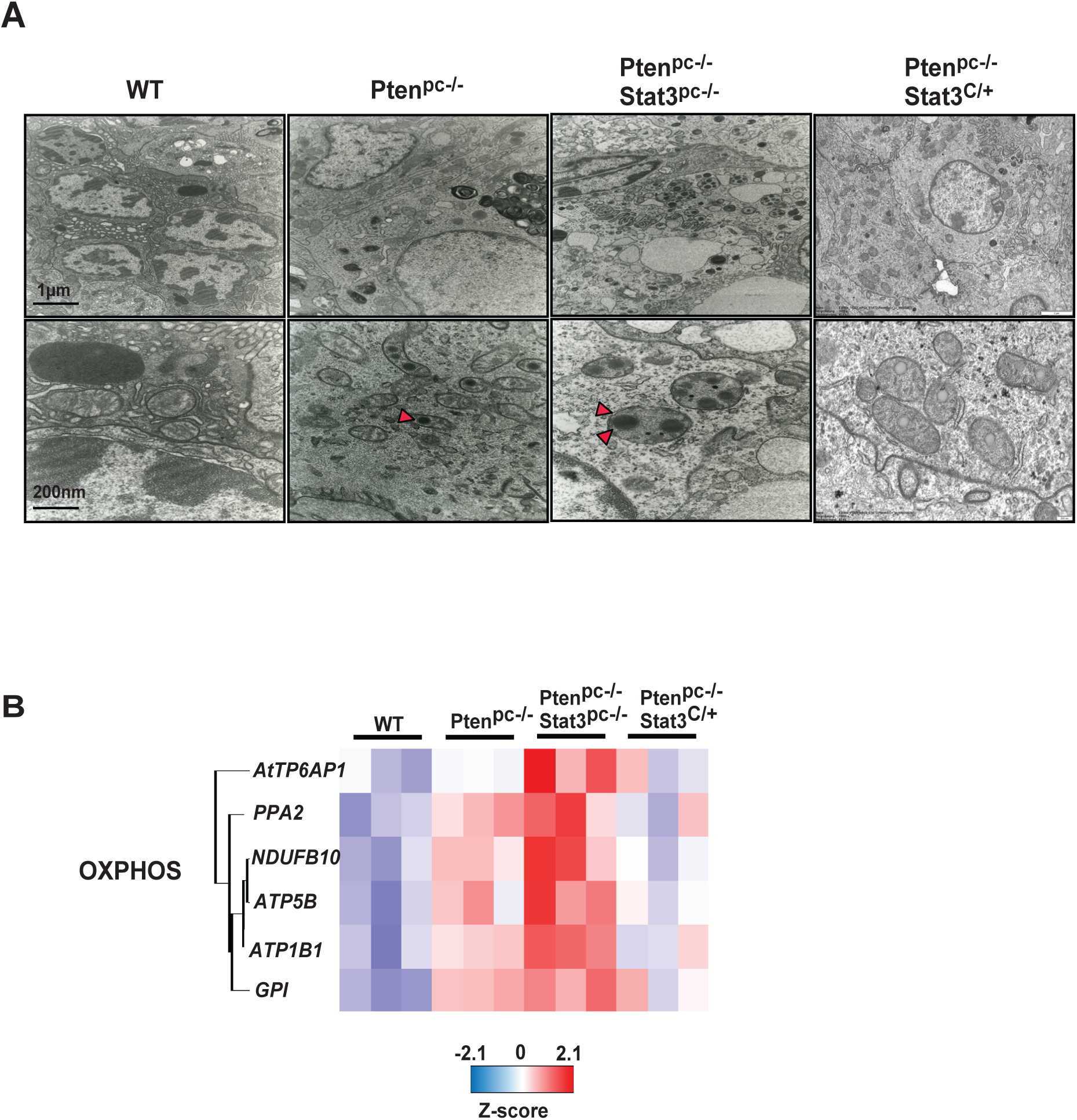
(**A**) Transmission electron microscopy (TEM) pictures of mitochondria from WT, *Pten^pc−/−^*, *Pten^pc−/−^Stat3^pc−/−^*and *Pten^pc−/−^Stat3^C/+^*prostates, 19 weeks of age mice. Black arrowheads show defected mitochondrial shape. Stat3 activation in *Pten^pc−/−^Stat3^C/+^* cells reversed defected mitochondrial phenotype. (**B**) The heatmap of murine PCa proteomics with significant enrichment of genes involved in regulation of oxidative phosphorylation. 19- week-old WT, *Pten^pc−/−^*, *Pten^pc−/−^Stat3^pc−/−^*and *Pten^pc−/−^Stat3^C/+^* mice (n=3).

**Supplementary Fig. 4.**
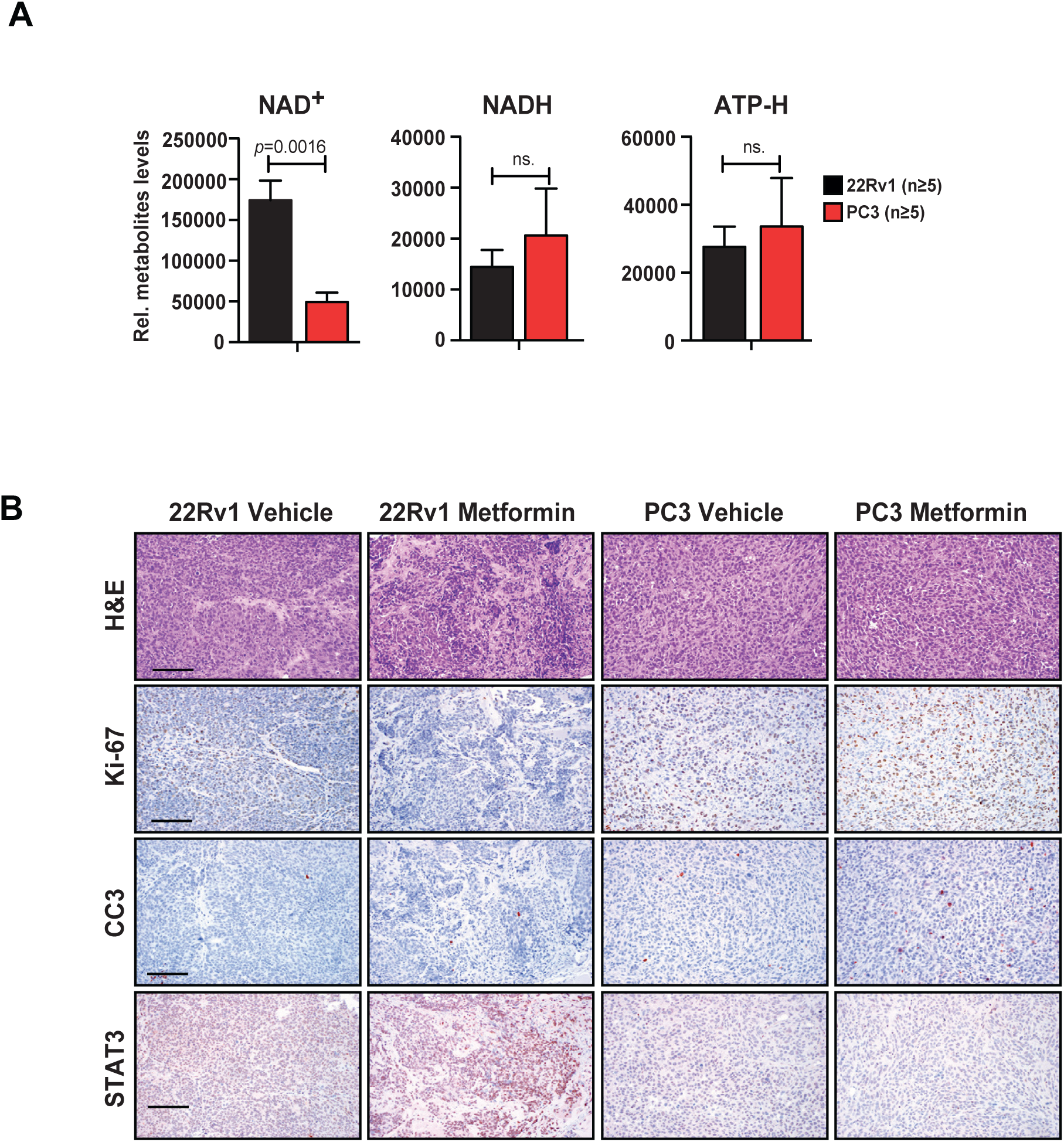
(**A**) HRSM measurement of NAD+, NADH and ATP in 22Rv1 and PC3 tumors (n=5). (**B**) H&E and IHC stainings of Ki-67, Cleaved Caspase 3 (CC3) and STAT3 expression in vehicle versus metformin-treated xenografted tumors (n=5), scale bar 50 µm.

**Supplementary Fig. 5.**
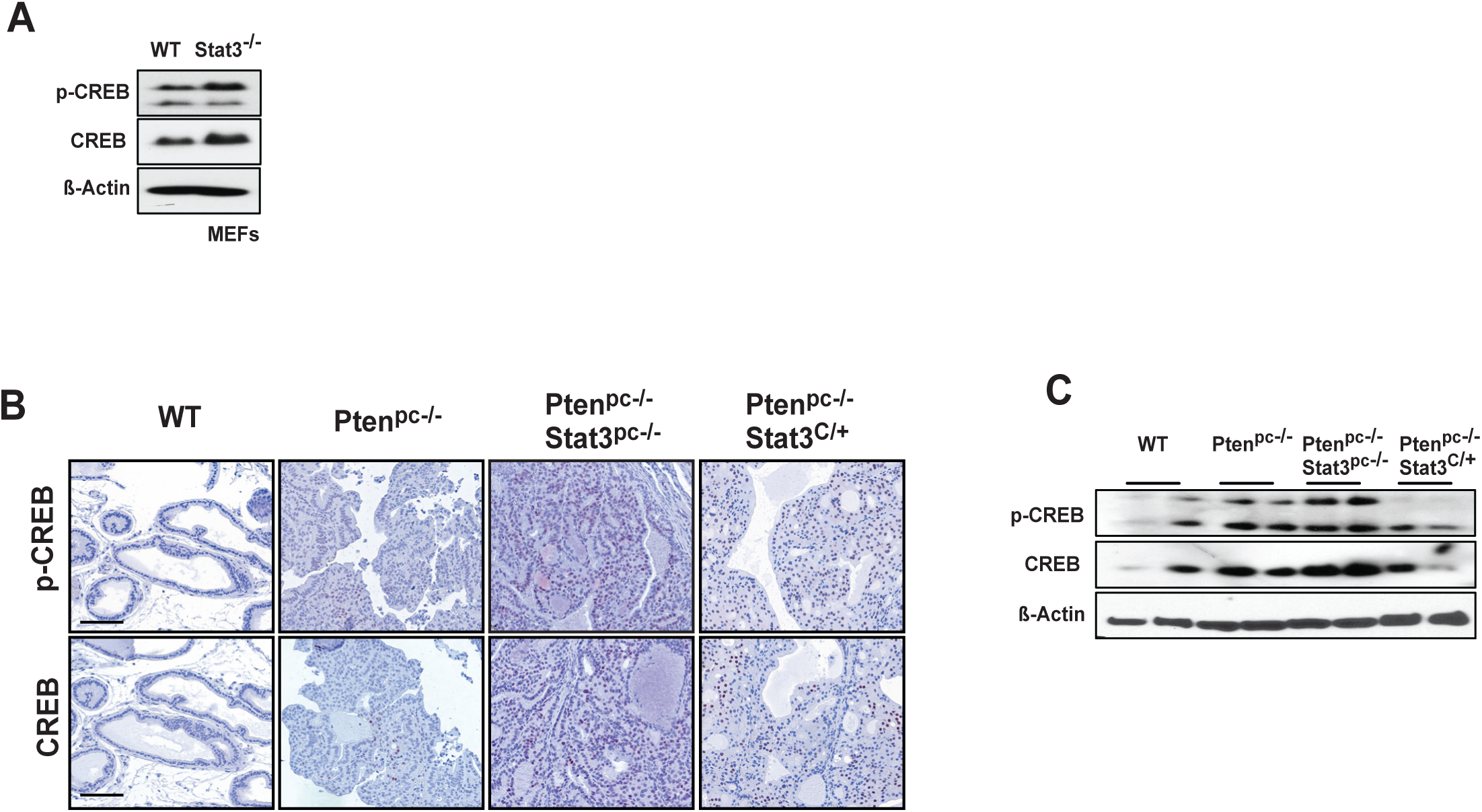
(A) Western blot analysis of p-CREB and CREB expression in WT and Stat3KO MEFs. β-actin serves as a loading control. (B) IHC analysis of p- CREB, and CREB in prostates from 19-week-old WT, Ptenpc−/−, Ptenpc−/− Stat3pc−/− and Ptenpc−/−Stat3C/+ mice. Scale bars, 100 μm. (C) WB analysis of p-CREB and CREB protein expression in prostates from 19- week-old WT, Ptenpc−/−, Ptenpc−/− Stat3pc−/− and Ptenpc−/−Stat3C/+ mice.

**Supplementary Fig. 6.**
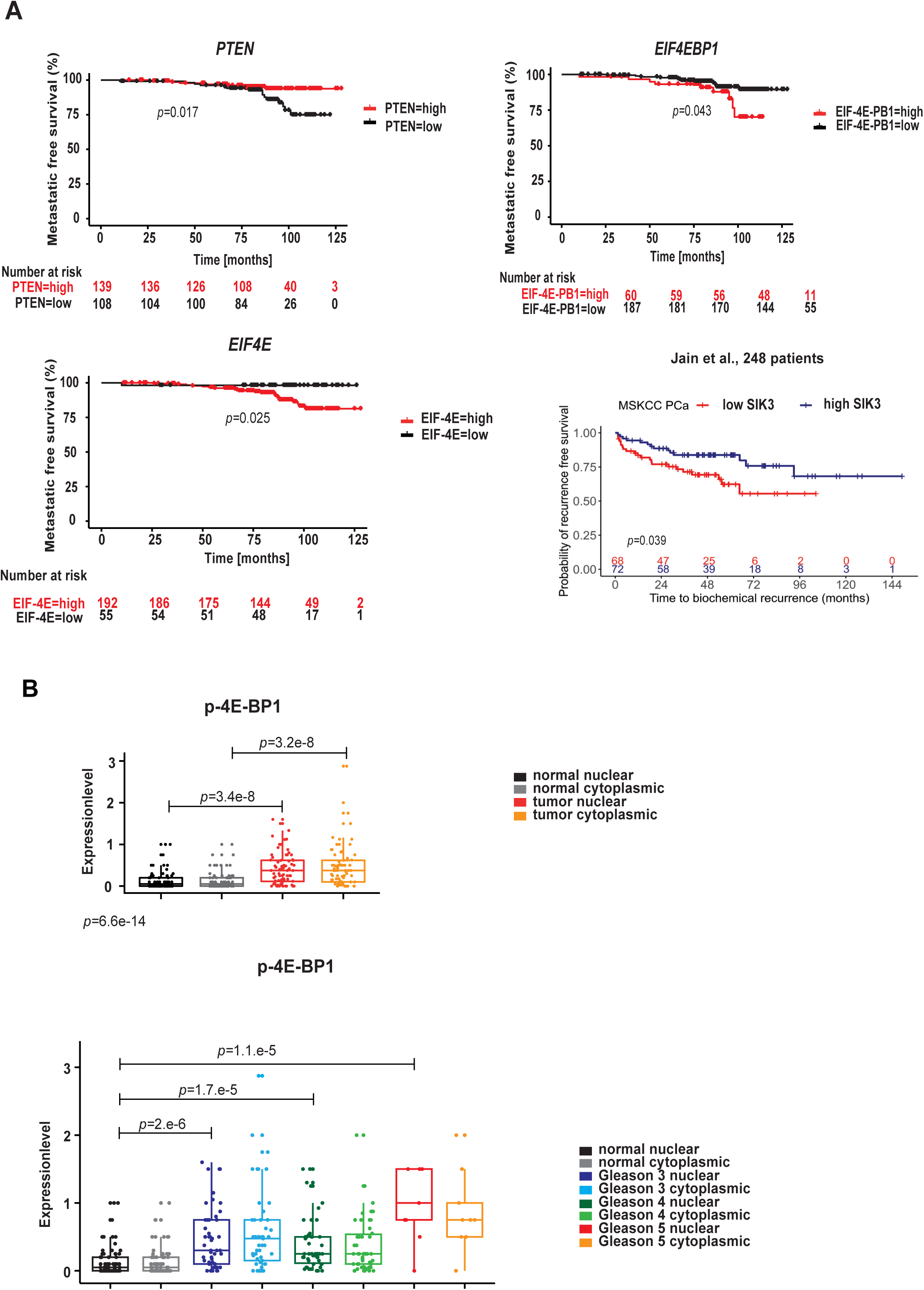
(**A**) *PTEN, EIF4EBP1, EIF4E mRNA* levels of 248 patients with localized/locally advanced PCa commencing radical radiotherapy (with/without ADT) analyzed by RNA-Seq (Jain et al., 2018)^55^ and Kaplan-Meier plot indicating time to biochemical recurrence in months for SIK3 mRNA in the MSKCC 218 PCa patients (Taylor et al., 2010)^38^. Groups were generated by a median split. Significance was estimated by log-rank test (95% confidence interval) and *p*-value was adjusted with Benjamini-Hochberg method. + = censored. (**B**) Boxplot representing protein expression of p-4E- BP in cytoplasmatic or nuclear stainings detected by IHC in normal-like glands or tumors in PCa patient TMAs (n=83). Evaluation and Gleason pattern annotations of nuclear and cytoplasmic IHC stainings of p4E-BP1 in PCa patient TMAs (n=83). Boxplots show median, 1st and 3rd quartiles, and whiskers extend to ± 1.5 interquartile range. Kruskal-Wallis and Dunn’s all pair tests were performed to assess significance (95% confidence interval).

**Supplementary Fig. 7.**
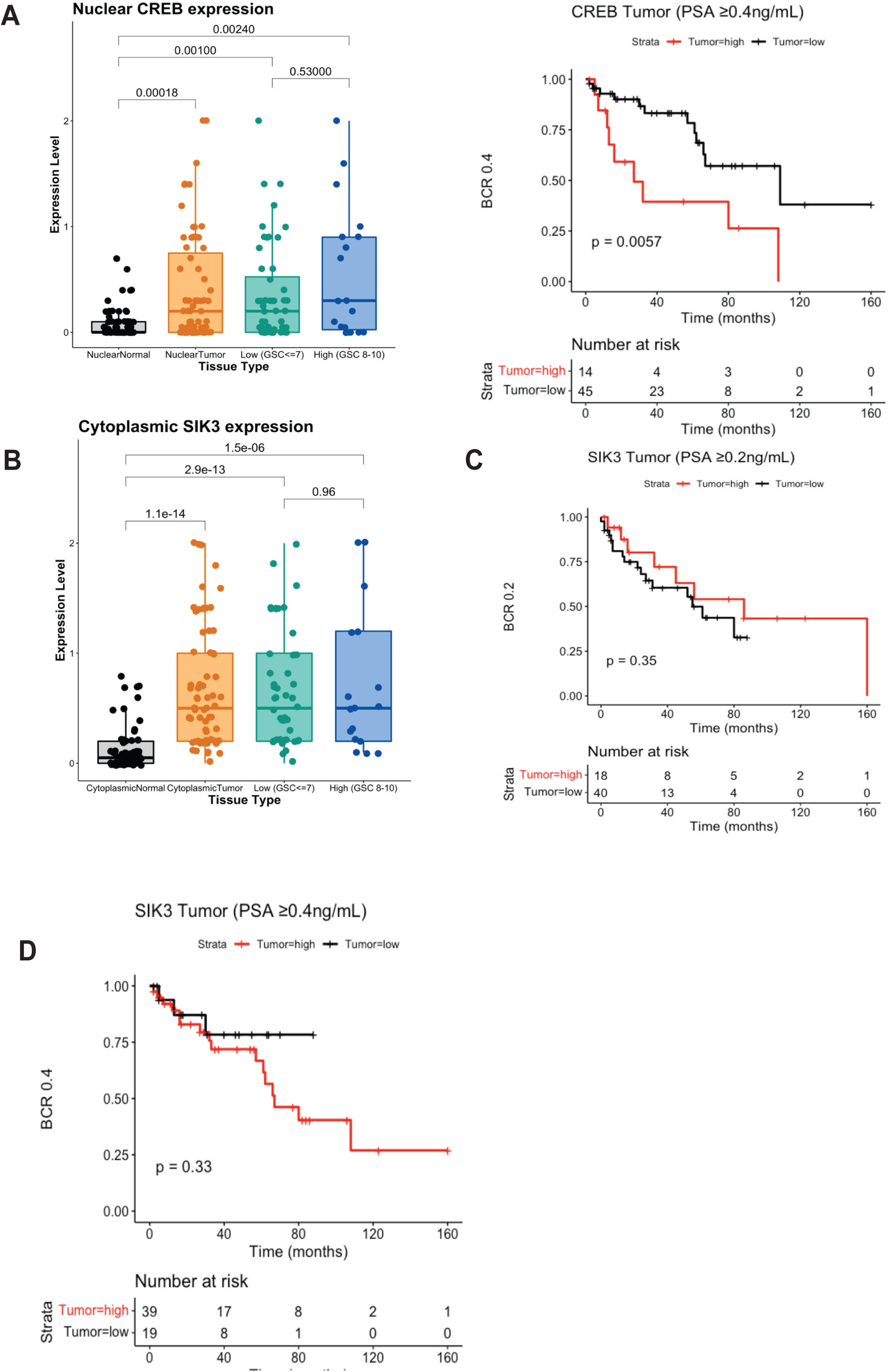
(**A**) Boxplot representing CREB nuclear protein expression stainings detected by IHC in PCa patient TMAs (n=83), (low GSS≤7 and high GSC=8-10). Kaplan–Meier analysis of BCR-free survival ratio based on CREB protein expression in a panel of 83 PCa patients (PSA ≥ 0.4ng/L). (**B**) Boxplots representing SIK3 cytoplasmic protein expression stainings detected by IHC in PCa patient TMAs (n=83), (low GSS≤7 and high GSC=8-10). **(C** and **D**) Kaplan–Meier analysis of BCR-free survival ratio based on SIK3 protein expression in a panel of 83 PCa patients (PSA≥0.2ng/L) and (PSA≥0.4ng/L).

**Table S1.**
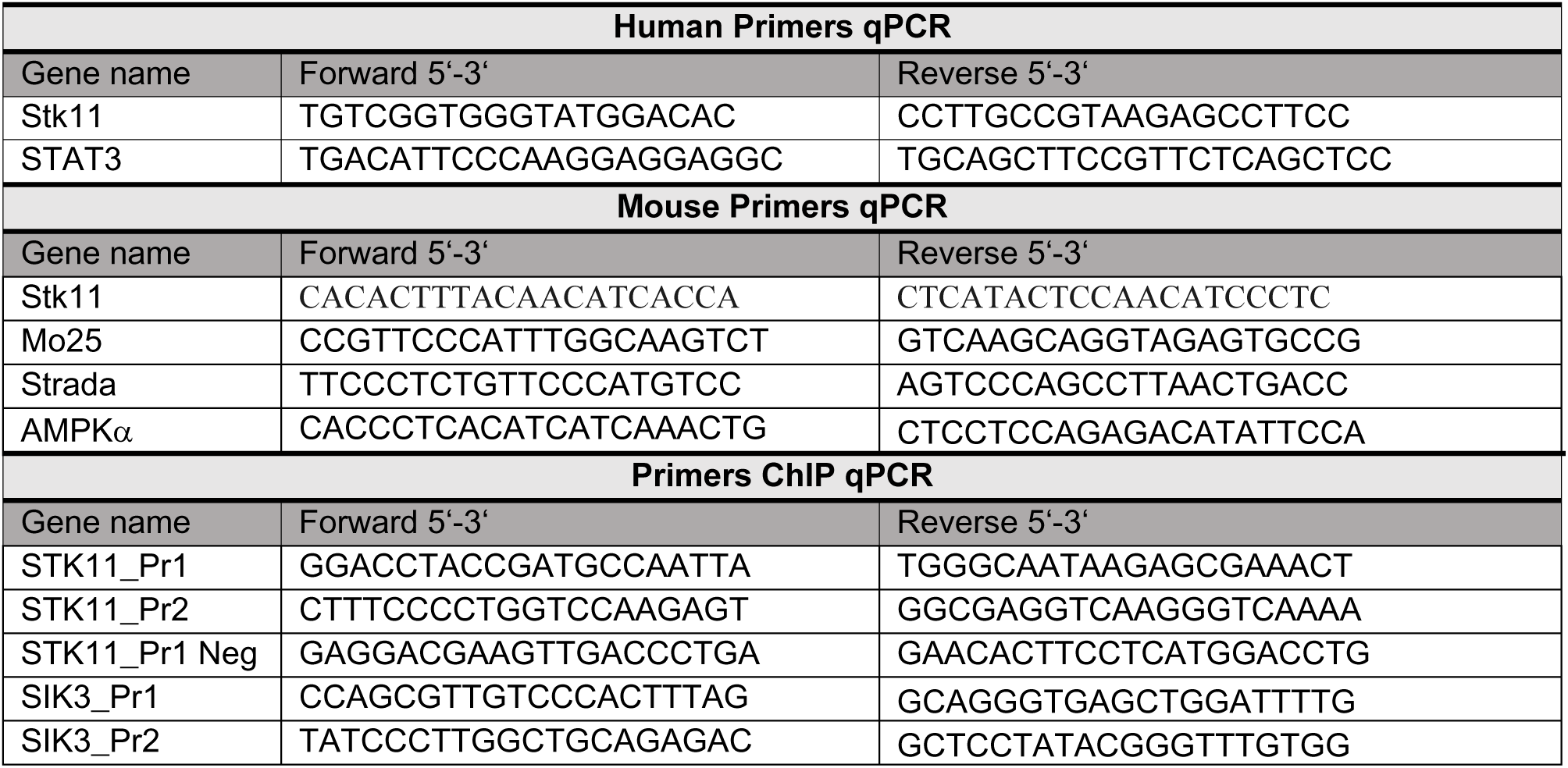

## Methods

### Generation of transgenic mice

The prostate epithelium-specific deletion was generated by the PB-Cre crossed with *Pten^loxP/loxP^* and/or Stat3*^loxP/loxP^* conditional mice as described previously(8). To generate constitutively activated *Stat3*, we took advantage of *Stat3^C/+^*(30) and crossed them with *Pten^loxP/loxP^* mice. All cohorts were in a C57BL/6 and 129/Sv mixed genetic background. Animal experiments were reviewed and approved by the Austrian ministry authorities and conducted according to relevant regulatory standards (BMWFW-66.009/0281-I/3b/2012 and BMWFW-66.009/0088-WF/V/3b/2018). The conditional tissue-specific prostate and lung metastatic samples with *Pten*^pc−/+^ (heterozygous) and/or *Lkb1* alterations were kindly provided by Arkaitz Carracedo and described previously by Hermanova et al.^34^.

### Clinical specimens

We retrospectively analyzed PCa patients diagnosed with both t2DM and PCa who underwent an open retropubic or robotic assisted (Da Vinci) RPE. In total, 570 tissue samples from 95 patients were collected. Cylindrical samples including three cancer areas and three benign areas were re-located from formalin-fixed, paraffin-embedded tissue blocks to the TMA block. Embedded hepatic cells as well as prostate cells (LNCaP and PC3) were used as controls. A tissue micro array (TMA) was assembled using a manual tissue arrayer (Beecher Instruments). The study was approved by the Ethical Committee of the Medical University Innsbruck (study number AN2014-0145 336/4.24) and written informed consent to participate in research studies was obtained from all patients.

A second TMA PCa cohort was obtained from the Department of Pathology of the Medical University of Vienna (MUW), Vienna, Austria. The FFPE material originated from 83 primary PCa patients and seven bladder cancer patients who underwent radical prostatectomy at the General Hospital of Vienna from 1993 to 2015. Use of patient FFPE material in this study was approved by the Research Ethics Committee of the Medical University Vienna, Austria (1877/2016). Staining intensities (Int) were rated from weak (= 1) to strong (= 3) and percentage (Perc) of positive cells were evaluated. Thereby, the overall expression level (EL) was derived:

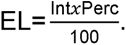

For group comparisons, Pearson chi-square normality test, Q–Q (quantile-quantile) plots and density plots were applied to test for normality and to visually inspect data. Levene’s Test was applied to test for homogeneity of variance. Either parametric (ANOVA) or non-parametric (Kruskal-Wallis) tests were used. Pairwise comparisons were performed using Bonferroni multiple comparisons of means after ANOVA and Dunn’s all pair test after Kruskal-Wallis test. Significance was defined as *p*-value ≤ 0.05. Statistical tests were performed using the R software environment with packages DescTools v.0.99.28, PMCMRplus v.1.4.1 and nortest v.1.0-4. Plots were generated with ggpubr_0.2 and ggplot2_3.3.0. Data were processed using tidyverse v.1.2.1.

### Plasma samples

The study was approved by the Ethics Committee of the Medical University of Graz (approval number 21-228 ex 09/10), conducted according to the Declaration of Helsinki, and written informed consent was obtained from all patients. At the time of the first blood collection, 6 (14.0%) patients were castration sensitive (CSPC) and 37 (86.0%) patients were CRPC. 95 plasma samples derived from 43 patients with metastatic PCa^79^. The majority of cases (35/43; 81.4%) displayed typical high-grade prostate adenocarcinoma features, five cases (11.7%) were poorly differentiated prostate cancers, one case each (2.3%) had undifferentiated or glandular histology and in one case (2.3%) the histology could not be obtained. No case showed neuroendocrine differentiation or exhibited small-cell neuroendocrine features. In the following are detailed histories of the patients with serial plasma DNA analyses.

### Immunohistochemistry and histological analysis

Immunohistochemistry and haematoxilin/eosin staining was performed with formalin-fixed paraffin-embedded (FFPE) tissue using standard protocols using consecutive sections. The following antibodies were used for immunohistochemistry: STAT3 (1:100 dilution; CST, #9139), LKB1 (1:100 dilution; Abcam; ab15095), p-AMPK 1:100 dilution; CST, #2535), p-4E-BP1 1:100 dilution; CST, #2855), 4E-BP1 1:100 dilution; CST, #9644), p-S6 1:100 dilution; CST, #2211), PTEN 1:100 dilution; CST, #9188),Ki-67 (1:1,000 dilution; Novocastra; NCL-KI-67-P) and Cleaved Caspase 3 (1:200 dilution; Cell Signaling, #9661). p-CREB 1:100 dilution; CST, #9198), CREB 1:100 dilution; CST, #9197), SIK3 (1:100 dilution; Novus Biologicals, NBP2-47278).

IHC was performed on a Discovery-XT staining device (Ventana). The following antibodies were used: AMACR/p63 immunohistochemistry (IHC) double staining (Monoclonal Rabbit Anti-Human AMACR, clone 13H4, Dako, Code M361601-2, 1:100, CC1 and Ventana Anti-p63 (4A4) Mouse Anti-Human Monoclonal, Catalog Number: 790-4509, BM, CC1).

For quantifying expression levels an established semi-quantitative “quick score” system combining the proportion of positive cells and the average staining intensity based on the method first described by Detre et al.^80^ was used. Briefly, quick score categories were based on both the proportion (denoted category A) and intensity (denoted category B) of positively stained cells. The proportion of positive cells (category A) was stratified into 4 groups (0: negative, 1: ≤30%, 2: 30–60%, 3: ≥60%). Average staining intensity (category B) corresponding to the presence of negative, weak, intermediate, and strong staining was given a score from 0 to 3, respectively. An average multiplicative quick score (category A × category B) for each TMA tissue core was subsequently obtained.

For electron microscopy, mouse prostate tissue was cut into 2 mm pieces and fixed in 1.6% glutaraldehyde overnight. Photographs were taken at a 4,000× magnification using a transmission electron microscope.

All images were taken with a Zeiss AxioImager Z1, and quantification was performed with HistoQuest (TissueGnostics GmbH, Vienna, Austria, www.tissuegnostics.com) as described in detail in Schlederer *et al*. (*81*). In brief, haematoxylin staining was used for cell identification. The range of intensities of the master marker (haematoxylin) and the immunohistochemical stainings were set by autodetection of the software. All images were analyzed with the identical settings after adjustments. The results are visualized in dot plot scattergrams and/or histograms. Cut-offs (to differentiate between positive and negative cells) and gates (to accentuate between cell populations) were set in the dot blots. For statistical analysis, the raw data were imported into GraphPad Prism 6 (GraphPad Software), analyzed for significance, and processed for data output. All images were taken with the same exposure time, signal amplification and objectives.

### Western blot analysis

For protein expression analysis by western blot, frozen tissue samples and cell lysates were prepared as described^8^. Blots were blocked with 5% BSA or 5% non-fat dry milk in 1 × TBS/0.1% Tween-20 for 1□h and incubated with the primary antibody overnight at 4□°C. Primary antibodies were reactive to STAT3 (1:1,000 dilution; CST #9139), LKB1 (1:1,000 dilution; CST #3047), p-4E-BP1 (1:1,000 dilution; CST #9451), 4E-BP1 (1:2,000 dilution; CST #9644), p-S6 (1:1,000 dilution; CST #4858), S6 (1:2,000 dilution; CST #2217), p-CREB (1:1,000 dilution; CST #9198), CREB (1:1,000 dilution; CST #9104), SIK3 (1:1,000 dilution; Novus Biologicals #47278), AR (1:1,000 dilution; CST #5153), PSA (1:1,000 dilution; CST #5365).

### RNA and qRT–PCR

Total RNA was isolated using Trizol (Invitrogen) according to the manufacturer’s instructions. For quantitative reverse transcription PCR (qRT– PCR) analysis, 1□μg of total RNA was reverse transcribed to cDNA using the Transcriptor First-Strand cDNA Synthesis kit (Fermentas). qRT–PCR was performed in triplicate with aa MxPro3000 and SYBR GreenERqPCR mix (Invitrogen). For qPCR analysis CFX96 Real-Time PCR Detection System (BioRad, Hercules, CA, USA) was employed using RNA expression of target genes relative to β-actin was quantified by 2ΔΔCT method. The relative amount of specific mRNA was normalized to β-actin in each sample. Primer pairs are listed in Supplementary Table 1.

### Cell culture

Primary WT and *Stat3*-null MEF were isolated by trypsin treatment of individual littermate E13.5 embryos from a cross of *Stat3^+/−^* heterozygous mice. *Stat3^+/−^* mice were generated from conditional *Stat3^+/fl^* mice^82^ by deletion of the conditional allele *in vivo* using *Mox2-Cre*. Cells were amplified and used in experiments starting at passage 2. *Stat3^−/−^* MEFs were grown in DMEM supplemented with 10% FBS, 2□mM L-Glutamine, 0.1□mM NEAA, 20□mM HEPES and pen/strep using standard techniques. For *in vitro* cultures LNCaP, RWPE-1 and PC3 were cultured under standard conditions.

Short hairpin-mediated knockdown was performed as previously described by Eberl *et al*.^83^. For the knockdown of *STAT3* in 22Rv1 cells, the following short hairpin RNA (shRNA) constructs from the Mission TRC shRNA library (Sigma) were used: scrambled control shRNA (SHC002), shSTAT3#456 (TRCN0000071456), shSTAT3#843 (TRCN0000020843), shLKB1#g1 (TRCN0000000408) and shLKB1#2 (TRCN0000195299). Transduced cells were selected for puromycin resistance, and the knockdown was verified via WB. The PC3 cells transfected with pcDNA3-TOPO-STAT3-V5 or empty vector (pcDNA3-TOPO, pc3.1 were described previously by Pencik *et al*. ^8^.

### IC50

5,000 cells were plated in 96 well flat bottom plates and treated in triplicates with DMSO (negative control), Bortezomib (10 µM; positive control), or compounds of interest. Cell viability was assessed after 72 h using the CellTiter Blue assay (Promega, Madison, WI, USA) according to the manufacturer’s protocol. IC_50_ values were determined using GraphPad Prism 9 by non-linear regression.

### Xenograft models in NSG mice

LNCAP cells were grown in RPMI with 10% FBS, 1% Pen/Strep. Cells were split 24h before harvesting cells were detached with trypsin, washed twice in PBS and counted. Cells were suspended in PBS at 2 x 10^6^/ml, mixed 1:1 with Matrigel (Corning), and kept on ice until injection. Nine weeks old NSG mice received sub-cutaneous injections of 200 μl tumor matrigel mix, so that each mouse received 1 x 10^6^ cells. On day 2 mice received either oral gavage containing Ruxolitinib dissolved in a PBS/DMSO solution (20% DMSO), at 50 mg/kg; or a control PBS/DMSO (Ruxolitinib n=5; Control n=5). Mice were treated every subsequent day for a total of 10 days. Mice were monitored for tumor development and sacrificed 24 days into the experiment when one of the tumors had exceeded the size limit of 1.2 cm in diameter. Tumors were dissected and weighed, then fixed in formalin (10%) for further analysis.

Cells (PC3 or 22Rv1) were harvested using trypsin, washed twice in PBS and counted. Cells were suspended in PBS at 2x10^6^/ml, mixed 1:1 with Matrigel (Corning), and kept on ice until injection. Nine-week-old NSG mice received subcutaneous injections of 200 µl tumor matrigel mix, so that each mouse received 1 x 10^6^ cells per flank. Mice were monitored for tumor development; once the tumors reached 100-150 mm^3^ in size (approx. three weeks after inoculation), mice were treated with vehicle (0.9% saline) or metformin (Sigma, PHR1084) reconstituted in 0.9% saline was administered via intraperitoneal injection once daily for 12 (PC3) and 19 (22Rv1) consecutive days, respectively. Tumors were dissected and weighed, then fixed in formalin (10%) for further analysis.

### Establishing serially transplantable AD and or AI PCa xenografts

AD (i.e., androgen-sensitive) xenograft tumors, LNCaP, were routinely maintained in intact immunodeficient NSG mice. To establish the CR lines, parental AD tumor cells were purified, mixed with matrigel, injected subcutaneously and serially passaged in surgically castrated immunodeficient mice. The complete method is described by Li *et al*.^56^.

### High-resolution mass spectrometry (HRMS)

The xenografts were extracted in 1 mL cold methanol. The extracts were dried in a vacuum concentrator and reconstituted in methanol to concentrations normalized to 40 mg xenograft tissue/100 µL methanol. The samples were analyzed with liquid chromatography electrospray-time-of-flight mass spectrometer (LC-ESI-ToF-MS) (maXis impact, Bruker Daltonics, Bremen, Germany). The mass spectrometer was operated with a capillary voltage of ±4 kV and a plate offset voltage of ±500 V. Nitrogen gas (200 °C) administered at 8 L/min was used as desolvation gas. The nebulizer pressure was 2 bar. The digitizer sample rate was set at 4 GHz and profile spectra were collected at a rate of 1 Hz. NAD+ was measured in positive mode and ATP and NADH were measured in negative mode. For the positive mode analysis, ten µL reconstituted extract was injected onto a Waters Atlantis HILIC Silica column (3 µm, 2.1 × 150 mm) (Waters, Milford, MA) (30 °C, flow rate was 0.25 mL/min). The aqueous mobile phase A consisted of 10 mmol/L ammonium formate (Sigma-Aldrich) with 0.1% (v:v) formic acid (Fisher chemicals, Hampton, NH). Mobile phase B consisted of acetonitrile (VWR Chemicals) with 0.1% formic acid (v:v). The following gradient, expressed as percentage of mobile phase B, was used: 0 min 95%, 0.5 min 95%, 10.5 min40%, 15 min 40%, 17 min 95%, and 32 min 95%. For the negative mode, ten µL reconstituted extract was injected onto Waters Xbridge BEH amide column (3.5 µm, 4.6 × 100 mm) (30 °C, flow rate was 0.25 mL/min). The aqueous mobile phase A consisted of 95:5 (v:v) water:acetonitrile supplemented with 20 mmol/L ammonium acetate and ammonium hydroxide (Sigma-Aldrich). Mobile phase B was 100% acetonitrile. The following gradient program, expressed as percentage of mobile phase B, was used: 0 min 85%, 3 min 30%, 12 min 2%, 15 min 2%, 16 min 85%, and 23 min 85%.

### Chromatin immunoprecipitation assays (ChIP)

Soluble chromatin preparation and ChIP assays were carried out as described previously (Hauser et al.)^84^ with some modifications. In short, cells were crosslinked with 1% v/v formaldehyde for 10 min at room temperature and the crosslink was stopped by the addition of glycine to a final concentration of 125 mM for 5 min while shaking. Chromatin was sonicated using a Twin Bioruptor (Diagenode) 30 s on/off for 15 cycles at 4°C. Two hundred microgram of chromatin was used for IP with 10 μl of STAT3 (1:50, Cat#12640, Cell Signaling) and 4 μg of IgG (1:250, Cat#10500C, Thermo Fisher Scientific) antibodies and incubated overnight. Protein–antibody complexes were bound to magnetic protein G beads (Life Technologies) for 4–5 h and washed with standard IP wash buffers for 10 min at 4°C. The crosslink was reversed by addition of 0.05 volume of 4M NaCl overnight at 65°C. After proteinase K digestion, DNA was recovered by phenol– chloroform–isoamylalcohol extraction and dissolved in 200 μl H2O. Real-time PCR of diluted ChIP DNA and corresponding input DNA was performed on ViiA 7 Real-Time PCR system (Thermo Fisher Scientific). Primer sequences used for ChIP are listed in Supplementary Table 1. Known STAT3 binding sites in BATF and JUNB promoters described in Tripathi et al.^85^ were chosen as positive controls and confirmed by extraction of corresponding peaks from ENCODE STAT3 ChIP-Seq HeLa-S3 data (ENCSR000EDC) with UCSC Genome Browser (http://genome.ucsc.edu). For the generation of STK11 and SIK3 primer pairs, a STAT3 binding site in the promoter region of STK11 and SIK3 detected by ENCODE STAT3 ChIP-Seq HeLa-S3 and STK11 used in Linher-Meville et al. ^35^. Primer pairs were created with Primer3web v4.1.0 software^86^.

### Liquid chromatography tandem mass spectrometry (LC**-**MS/MS)

For LC-MS/MS proteomics, FFPE tumor material was used from 19-week-old WT, *Pten^pc−/−^*, *Pten^pc−/−^ Stat3^pc−/−^* and *Pten^pc−/−^Stat3^C/+^*mice (n*=*3). Blocks were sliced into 3-μm-thick sections, mounted on slides, and stained with hematoxylin and eosin. Tumor areas were marked by a pathologist. To obtain proteomic profiles solely from the tumor, stroma and immune cells were excluded from dissection. One hundred nanoliter (10 mm^2^ of 10 μm slides) of FFPE material per sample was used for analysis. Lysis of microdissected tissue was carried out in 50% trifluoroethanol (TFE), 5 mM dithiothreitol (DTT), 25 mM ammonium bicarbonate (ABC) at 99°C for 45 min. followed by 5-min. sonication (Bioruptor, Diagenode). After centrifugation at 16,000 g for 10 min., the cleared protein lysate was alkylated with 20 mM iodoacetamide for 30 min. at room temperature. Upon vacuum centrifugation, digestion was carried out in 5% TFE, 50 mM ABC to which 0.15 μg of LysC and 0.15 μg of trypsin were added for digestion overnight at 37°C. The following day, digestion was arrested by adding trifluoroacetic acid (TFA) to 1% and the digestion buffer removed by vacuum centrifugation. Peptides were suspended in 2% acetonitrile and 0.1% TFA and purified on C18 StageTips. Finally, purified peptides were resolved in 2% acetonitrile and 0.1% TFA, and the entire sample was injected for MS analysis in a single-shot measurement. Protocols were adapted from Roulhac et al.^87^ and Wang et al.^88^.

LC-MS/MS analysis was performed on an EASY-nLC 1000 system (Thermo Fisher Scientific) coupled online to a Q Exactive HF mass spectrometer (Thermo Fisher Scientific) with a nanoelectrospray ion source (Thermo Fisher Scientific). Peptides were loaded in buffer A (0.1% formic acid) into a 50-cm-long, 75-μm inner diameter column in house packed with ReproSil-Pur C18-AQ 1.9□μm resin (Dr. Maisch HPLC GmbH) and separated over a 270-min gradient of 2–60% buffer B (80% acetonitrile, 0.1% formic acid) at a 250 nl/min flow rate. The Q Exactive HF operated in a data-dependent mode with full MS scans (range 300–1,650 *m*/*z*, resolution 60,000 at 200 *m*/*z*, maximum injection time 20 ms, AGC target value 3e6) followed by high-energy collisional dissociation (HCD) fragmentation of the five most abundant ions with charge ≥ 2 (isolation window 1.4 *m*/*z*, resolution 15,000 at 200 *m*/*z*, maximum injection time 120 ms, AGC target value 1e5). Dynamic exclusion was set to 20 s to avoid repeated sequencing. Data were acquired with the Xcalibur software (Thermo Scientific). Xcalibur raw files were processed using the MaxQuant software v.1.5.5.2 (Cox & Mann, 2008)^89^, employing the integrated Andromeda search engine (Cox et al., 2011b)^90^ to identify peptides and proteins with a false discovery rate of < 1%. Searches were performed against the Mouse UniProt database (August 2015), with the enzyme specificity set as “Trypsin/P” and 7 as the minimum length required for peptide identification. N-terminal protein acetylation and methionine oxidation were set as variable modifications, while cysteine carbamidomethylation was set as a fixed modification. Matching between runs was enabled to transfer identifications across runs, based on mass and normalized retention times, with a matching time window of 0.7 min. Label-free protein quantification (LFQ) was performed with the MaxLFQ algorithm^91^ where a minimum peptide ratio count of 1 was required for quantification. Data pre-processing was conducted with Perseus software, v.1.5.5.5 for mouse data. Data were filtered by removing proteins only identified by site, reverse peptides, and potential contaminants. After log2 transformation, biological replicates were grouped. Label-free protein quantification intensities were filtered for valid values with a minimum of 70% valid values per group, after which missing data points were replaced by imputation. The resulting data sets were exported for further statistical analyses using R. Filtered, normalized, and log2-transformed data were imported, and PCA and unsupervised hierarchical clustering were performed. Plots were generated with ggplot2 v.3.1.1. (Wickham, 2016)^92^, gplots v.3.0.1.1 and EnhancedVolcano v.1.0.1 R packages.

### Statistical analyses

Data were analyzed using GraphPad Prism 6 software. For comparing two groups unpaired Students *t*-test and for comparing more than two groups Tukey’s *post hoc* test was used. Fisher’s exact test was employed when differences in distributions within groups were monitored. For Kaplan–Meier analysis and log-rank statistical evaluation of time to BCR, as well as evaluation of prognostic power in univariate and multivariate analysis, we used the IBM SPSS version 22 program. Survival analyses were performed using R packages survival_3.2-7 and survminer v0.4.6 Univariate Cox proportional hazards (PH) models were fitted for candidate genes. Log-rank tests for Cox PH significant genes with adj. *p*-value ≤ 0.01 were performed after a median split of samples by gene expression. All statistical tests were considered

### Software environment

Data acquisition, differential expression analyses, gene set testing and statistical analyses on RNA-seq data and tissue micro arrays were performed using the R software environment (https://cran.r-project.org/) with R versions 3.6.1. and 3.6.3 and packages as mentioned in the respective sections.

### Data acquisition

TCGA PRAD RNA-seq harmonized data (https://portal.gdc.cancer.gov/projects/TCGA-PRAD)(The Cancer Genome Atlas Research Network, 2017) were acquired by TCGAbiolinks v.2.10.5 (Colaprico *et al*., 2015)^93^. Data were normalized and transformed with edgeR v3.24.3. Normalized log2 counts of MSKCC PCa mRNA data (GEO: GSE21032) were derived from http://cbio.mskcc.org/cancergenomics/prostate/data/.

### Gene set testing

From TCGA PRAD data, a low-STAT3 (n=100) and a high-STAT3 (n=100) subset were selected, consisting of the 0.2^nd^ and the 1- 0.8^th^ quantile of overall STAT3 expression (cpm), respectively. After differential expression analysis (min. log-fold change = 0, max. *p*-value = 1) between low-STAT3 and high-STAT3 using limma v.3.40.6, genes were ranked by their moderated t-statistic. Gene set testing with fgsea_1.10.1) was performed on ranked genes with 10.000 permutations, a minimum gene set size =1 and an infinite maximum gene set size. *P*-values were adjusted by Benjamini-Hochberg correction. Significance was defined by an adj. *p*-value ≤ 0.05. Molecular Signatures Database (MSigDB) C2 gene sets were acquired by msigdbr_7.0.1 package.

### Oncomine database analysis

Gene expression analysis of *STAT3, PTEN, EIF4EBP1* and *SIK3* was performed in various prostate cancer datasets representing normal, tumor or metastatic samples deposited in the Oncomine Research Premium Edition database (Thermo Fisher, Ann Arbor, MI). For the analysis, the *P* value threshold was set to .05, the fold-change threshold was set to 1.5 and the gene rank threshold was set to “all.”

### Biochemical and metastatic recurrence analysis in PCa patients

The Walker et al.^94^ cohort consists of 322 FFPE prostatectomy samples. The Jain *et al*.^55^ cohort consists of 248 FFPE biopsy samples. The Walker et al cohort was dichotomised by median CREB1 expression. Cox proportional hazards regression method was used to estimate the univariate hazard ratio (HR) of the CREB1 expression categories. The relationship between CREB1 and AR expression was investigated in the Walker et al. and Jain et al. cohorts using the nonparametric Spearman’s rank-order correlation coefficient.

